# Jag1 Insufficiency Disrupts Neonatal T Cell Differentiation and Impairs Hepatocyte Maturation, Leading to Altered Liver Fibrosis

**DOI:** 10.1101/2022.10.24.513578

**Authors:** Jan Mašek, Iva Filipovic, Noémi Van Hul, Lenka Belicová, Markéta Jiroušková, Daniel V. Oliveira, Anna Maria Frontino, Simona Hankeova, Jingyan He, Fabio Turetti, Afshan Iqbal, Igor Červenka, Lenka Sarnová, Elisabeth Verboven, Tomáš Brabec, Niklas K. Björkström, Martin Gregor, Jan Dobeš, Emma R. Andersson

## Abstract

Fibrosis is a physiological tissue repair mechanism, but excessive fibrosis can disrupt organ function. Alagille syndrome (ALGS), which is caused by mutations in the Notch ligand *JAGGED1*, results in bile duct paucity, neonatal cholestasis, and a characteristic fibrotic response. Here, we show that *Jag1^Ndr/Ndr^* mice, a model for ALGS, recapitulates ALGS-like pericellular fibrosis. Single-cell RNA-seq and multi-color flow cytometry characterization of the liver and spleen revealed immature hepatocytes and paradoxically low intrahepatic T cell infiltration in cholestatic *Jag1^Ndr/Ndr^* mice, despite an enrichment in extrahepatic (thymic and splenic) regulatory T cells (Tregs). *Jag1^Ndr/Ndr^* lymphocyte immune and fibrotic capacity was tested with adoptive immune cell transplantation into *Rag1*^-/-^ mice, challenged with dextran sulfate sodium (DSS) or bile duct ligation (BDL). Transplanted *Jag1^Ndr/Ndr^* lymphocytes were less inflammatory with fewer activated T cells than *Jag1^+/+^* lymphocytes, in response to DSS. Cholestasis induced by BDL in *Rag1^-/-^* mice with *Jag1^Ndr/Ndr^* lymphocytes resulted in periportal Treg accumulation and three-fold less periportal fibrosis than in *Rag1^-/-^* mice with *Jag1^+/+^* lymphocytes. Finally, we show that the *Jag1^Ndr/Ndr^* hepatocyte expression profile and Treg overrepresentation are corroborated by transcriptomic data from children with ALGS. In sum, these data lead to a model in which Jag1-driven developmental hepatic and immune defects interact to determine the fibrotic process in ALGS.

## INTRODUCTION

Although fibrosis is a physiological tissue repair mechanism, excessive fibrosis is pathological and 45% of all deaths in the industrialized world are attributed to fibrosis (Henderson *et al*, 2020). Pathological liver fibrosis occurs in response to liver damage-induced chronic inflammation (Dobie *et al*, 2019). Hepatic fibrosis presents distinct fibrotic patterns including periportal, bridging or pericellular fibrosis, depending on the nature of the insult. Intriguingly, although Alagille syndrome (OMIM 118450; ALGS (Alagille *et al*, 1975)) and biliary atresia are both pediatric cholangiopathies, the associated liver repair and fibrotic mechanisms are distinct, with less bridging fibrosis but more pericellular fibrosis in ALGS than in biliary atresia (Fabris *et al*, 2007). One quarter of children with ALGS survive to adulthood without a liver transplant, and in many of these children serum bilirubin, cholesterol, and GGT, as well as pruritus and xanthomas, spontaneously improve (Kamath *et al*, 2020). Understanding the fibrotic process in ALGS could reveal disease-specific mechanisms and leverageable pathways to modulate specific aspects of liver fibrosis.

Notch signaling controls development and homeostasis, and mutations in the Notch signaling pathway leads to both developmental diseases and cancer (Mašek & Andersson, 2017; Siebel & Lendahl, 2017). ALGS is mainly caused by mutations in the Notch ligand *JAGGED1* (*JAG1,* 94%)(Mašek & Andersson, 2017; Oda *et al*, 1997), resulting in bile duct paucity and cholestasis, but immune dysregulation has also been described (Tilib Shamoun *et al*, 2015). Cholestasis-induced inflammation results in fibrosis and cirrhosis driven by T cells (Zhang & Zhang, 2020; Shivakumar *et al*, 2007), natural killer (NK) cells (Shivakumar *et al*, 2009), and myeloid cells (Jin *et al*, 2019; Henderson *et al*, 2020), attracted to the liver by injured liver parenchyma (Allen *et al*, 2011). Conversely, fibrosis is attenuated by regulatory T cells (Zhang & Zhang, 2020). While Notch regulates T cell lineage specification in liver and thymus (Radtke *et al*, 2004; Herman *et al*, 2005; Chen *et al*, 2019), the function of Jag1 in this process is less clear. Importantly, there is a ∼30% increase in infections in patients with ALGS (Tilib Shamoun *et al*, 2015)LJ, although this has been attributed to a Notch receptor-independent mechanism regulating T cell function itself (Le Friec *et al*, 2012) and it is unknown whether and how immune dysregulation in ALGS impacts liver disease and fibrotic progression.

Here, we investigated liver-immune system interactions in inflammation and fibrosis using *Jag1^Ndr/Ndr^* mice, a model of ALGS (Andersson *et al*, 2018), with orthogonal validation of corresponding transcriptomic signatures in liver biopsies from children with ALGS. Using multiomics across liver, spleen, and thymus, as well as adaptive immune transfer into lymphodeficient mice, we assess the lymphocytic capacity for interaction with liver cells. Our data demonstrate a multilayered role of Jag1 in the fibrotic process, affecting both hepatocyte maturation and injury response, as well as T cell differentiation and pro-fibrotic activity, providing a mechanistic framework for therapeutic modulation of Notch signaling in liver disease.

## RESULTS

### Jag1^Ndr/Ndr^ mice recapitulate ALGS-like pericellular fibrosis and portal hypertension

Fabris and colleagues have previously compared fibrotic architecture in ALGS and biliary atresia, demonstrating that patients with ALGS have less extensive bridging fibrosis but more pericellular fibrosis than patients with biliary atresia (Fabris *et al*, 2007). Therefore, we first asked whether *Jag1^Ndr/Ndr^* mice recapitulate ALGS-like pericellular fibrosis (Fabris *et al*, 2007) and portal hypertension (Kamath *et al*, 2020). *Jag1^Ndr/Ndr^* mice are jaundiced by postnatal day 3 (P3), with overt bile duct paucity and high levels of bilirubin at P10, which resolve by adulthood in surviving mice (Andersson *et al*, 2018).

Sirius red staining (SR) at P10, P30 and 3 months demonstrated an onset of heterogenous fibrosis at P10 with pericellular “chicken-wire” fibrosis, perisinusoidal fibrosis and denser fibrotic loci with pronounced immune cell infiltrate, which was notable in sections but not significantly enriched when quantified (**Fig. 1A-D**). In contrast, fibrosis was significantly elevated at P30, with occasional faint bridging fibrosis and pericellular fibrosis (**Fig. 1B,D**). At 3 months, fibrosis had partially resolved into sparse pericellular collagen strands (**Fig. 1C,D**). Splenic size is a common indicator of portal hypertension liver disease (Kamath *et al*, 2020). At birth, the *Jag1^Ndr/Ndr^* spleen was proportional to its body size, but splenomegaly developed by P10 and was further exacerbated by P30 (**Fig. 1E,F**). *Jag1^Ndr/Ndr^* mice thus mimic ALGS-like perisinusoidal, pericellular, and immune-associated fibrosis (Fabris *et al*, 2007) and develop splenomegaly, like patients with ALGS (Kamath *et al*, 2020; Vandriel *et al*, 2022). The atypical fibrotic reaction despite severe bile duct paucity and cholestasis (Andersson *et al*, 2018) mimics ALGS and shows that *Jag1* mutation alters the profibrotic mechanisms, which we next investigated.

**Figure 1:**
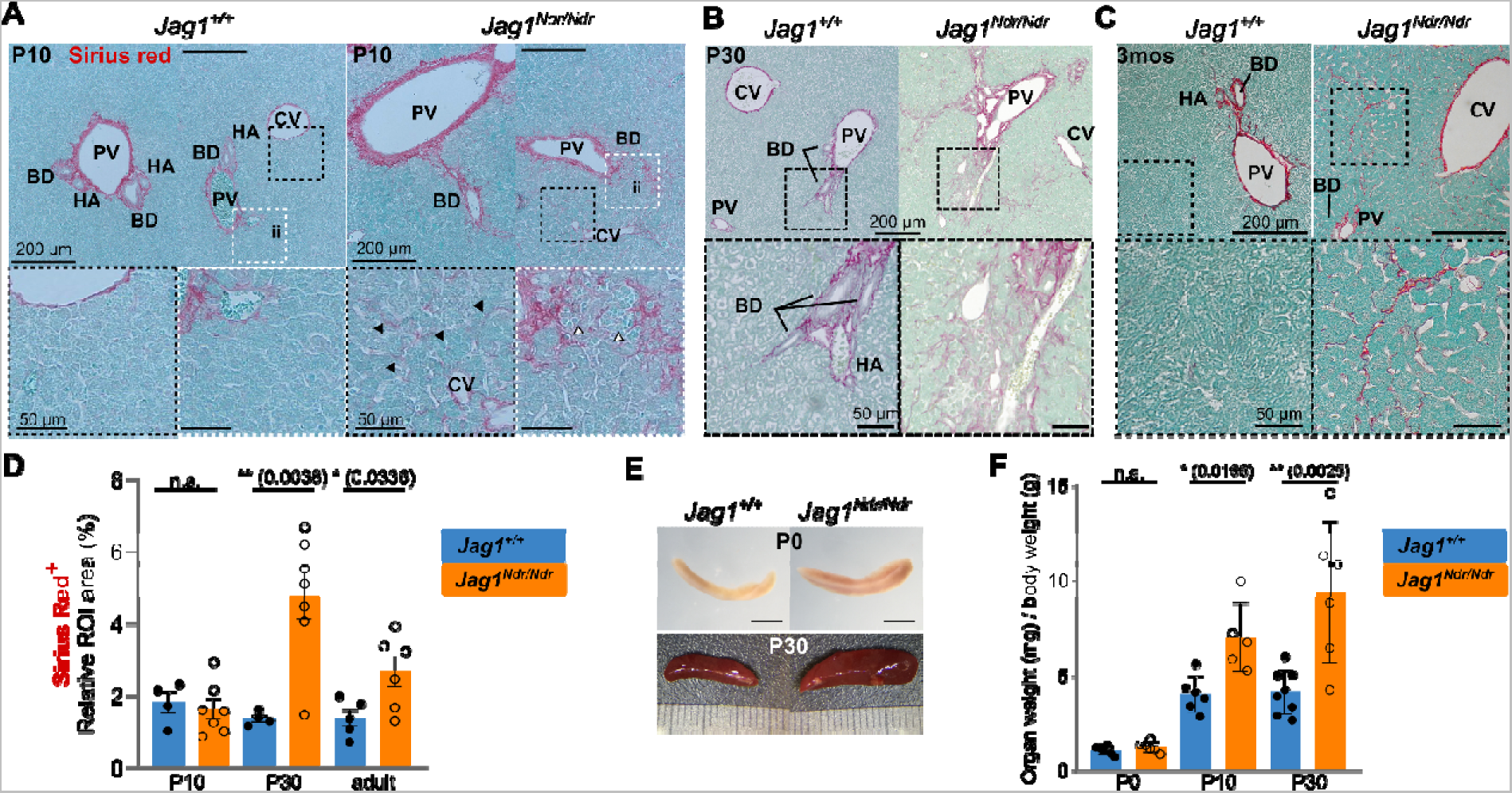
*Jag1^Ndr/Ndr^* mice display pericellular fibrosis and signs of portal hypertension. (**A-C**) H&E and Sirius red staining at P10 (A), P30 (B) and 3 months (C) in *Jag1^+/+^* and *Jag^Ndr/Ndr^* livers, with magnified regions boxed. Black arrowheads indicate pericellular and perisinusoidal fibrosis, white arrowheads indicate immune infiltration. (**D**) quantification of the Sirius Red^+^ at P10, P30 and 3 months in *Jag1^+/+^* and *Jag^Ndr/Ndr^* LLL liver sections. (**E,F**) Images (E) and weights (F) of spleens from *Jag1^+/+^* and *Jag^Ndr/Ndr^* mice at the indicated ages. The scalebar in E is 1mm. (Each data point represents a biological replicate. Mean ± SD, Unpaired, two-tailed Student’s t-test, p ≤ 0.05 = *, p ≤ 0.01 = **, n.s. not significant; LLL, left lateral lobe; BD, bile duct; PV, portal vein; CV, central vein; HA, hepatic artery; ii, immune infiltrate.

### Hepatocytes are less differentiated in Jag1^Ndr/Ndr^ mice and in Alagille syndrome, with dampened pro-inflammatory activation

To investigate which liver cell populations are affected by *Jag1* loss of function at the onset of cholestasis, we first analyzed *Jag1^+/+^*, *Jag1^Ndr/+^*, and *Jag1^Ndr/Ndr^* livers with single cell RNA sequencing (scRNA seq) at embryonic day (E) 16.5 and postnatal day 3 (P3) (**Fig. 2A**). 183,542 cells passed the quality control and filtering (**Fig. S1A,B**), and could be assigned to 20 cell clusters of parenchymal and non-parenchymal cells including tissue-resident Kupffer cells, and hematopoietic stem cells/erythroid-myeloid progenitors (HSc/EMP) and their derivatives: i) the myeloid lineage, ii) the lymphoid lineage, and iii) the erythroid lineage (Popescu *et al*, 2019; Liang *et al*, 2022) (**Fig. 2A, S1C-E**). The adult hepatocyte injury response program depends on HSC activation of NOTCH2 on hepatocytes, mediated by JAG1 (Nakano *et al*, 2017). We thus subset the 3,470 hepatoblasts and hepatocytes to analyze these in detail (**Fig. 2B,C, S2A**). While there were few or no significantly dysregulated genes between genotypes (**ST1-5**), there was an overt shift in cell population distributions between the genotypes (**Fig. 2D**). While hepatoblast and proliferating hepatoblasts ratios were similar across genotypes at E16.5, *Jag1^Ndr/Ndr^* P3 livers contained proportionally 3-times more Proliferative_Hepatoblasts than *Jag1^+/+^*, *Jag1^Ndr/+^* controls (31.5% vs 9.1%, 10.1%, respectively), but 1/3 less bone fide Hepatocytes (68.8% vs 90.1%, 89.2%, respectively), (**Fig. 2D, S2B**), suggesting a hepatocyte differentiation/maturation defect in *Jag1^Ndr/Ndr^* mice.

**Figure 2:**
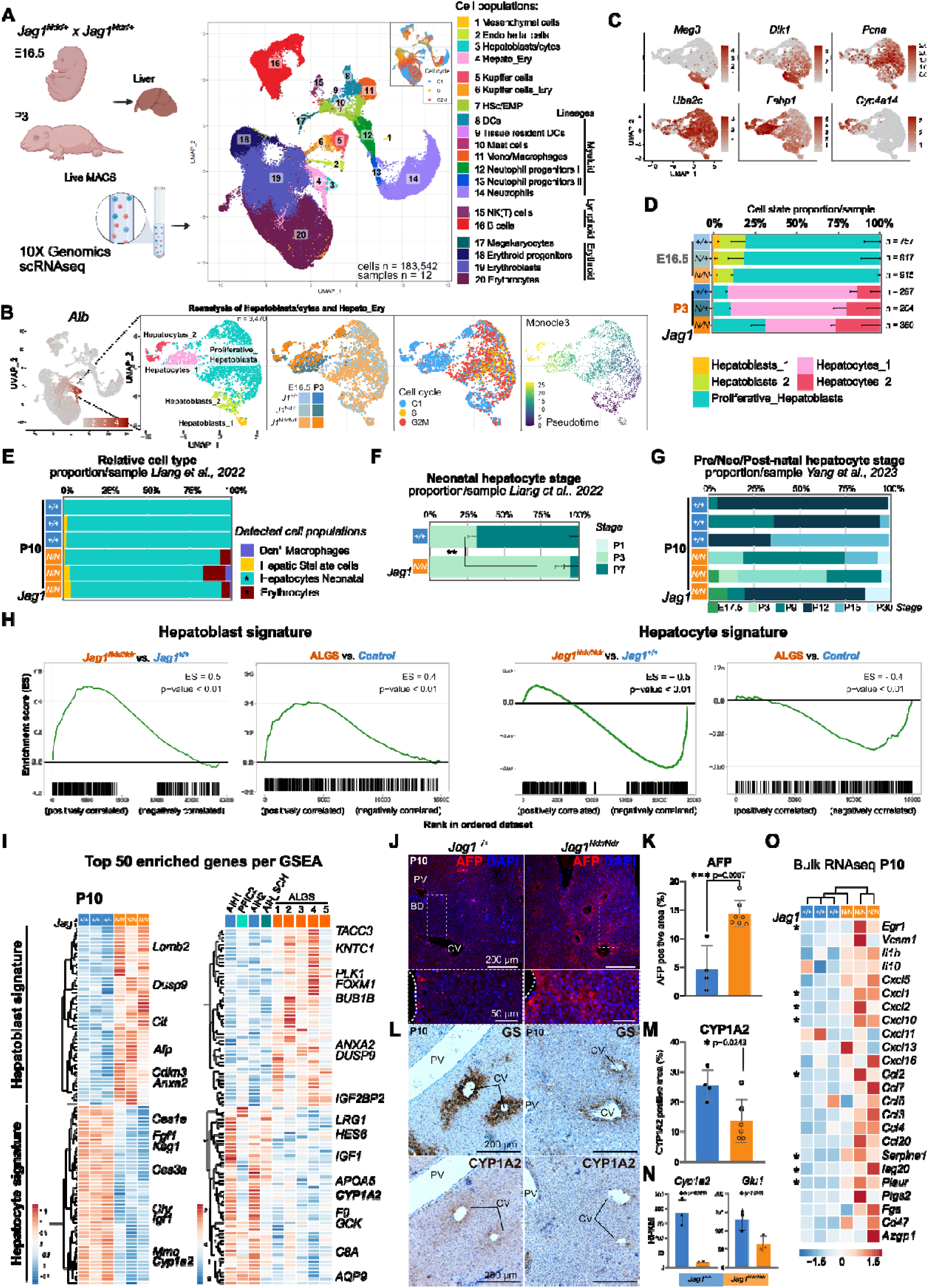
Injury-induced immature hepatocytes appear in neonatal liver of Jag1^Ndr/Ndr^ mice and patients with ALGS. (**A**) UMAP projection of 183,542 sequenced cells in cell type-annotated clusters, and cell cycle phase. Livers were collected at E16.5 or P3. Organ single-cell suspensions were analyzed by 10xGenomics scRNA sequencing. (**B**) UMAP of the Hepatoblast/cytes and Hepato_Ery subsets, reflecting their subclusters, genotypes, developmental stage, cell cycle status, and differentiation trajectory. (**C**) Feature plots with hepatoblast (*Meg3*, *Dlk1*), proliferation (*Pcna,Ube2c*), and hepatocyte (*Fabp1*, *Cyp4a14*) marker mRNA expression. (**D**) Average proportion of hepatic cell types per stage and genotype. (**E-G**) MuSiC deconvolution of P10 *Jag^Ndr/Ndr^* and Jag1^+/+^ whole liver bulk RNA seq using scRNAseq refence datasets (E), and stage-specific hepatocyte expression profile (F,G). (**H**) GSEA for hepatoblast and hepatocyte signatures in *Jag1^Ndr/Ndr^* vs *Jag1^+/+^* mice (top) or in patients with ALGS vs other liver diseases (bottom). (**I**) Top 50 enriched genes from each GSEA in (H) for mouse (left) and human (right) liver. (**J,K**) Staining for AFP and DAPI at P10 in *Jag1^+/+^* and *Jag1^Ndr/Ndr^* livers (J), with quantification (K). (**L-N**) Staining for GS and CYP1A2, with H&E counterstain (L), in consecutive sections at P10 in *Jag1^+/+^* and *Jag1^Ndr/Ndr^* livers, with quantification of CYP1A2 protein (M), and P10 whole liver *Cyp1a2* and *Glu1* mRNA expression (N). (**O**) Heat map of Egr1-induced proinflammatory genes expressed by hepatocytes in *Jag1^Ndr/Ndr^* and *Jag1^+/+^* mice at P10. (AIH = autoimmune hepatitis; PFIC2 = progressive familial intrahepatic cholestasis type 2; SCH = sclerosing cholangitis; (mean ± SD, Statistical analysis was performed by unpaired, two-tailed Student’s t-test, *, p <0.01, **, p <0.001), the rest (*, p_adj_ <0.01, Log2 FC>0.5).

To test whether hepatocyte differentiation was similarly delayed at later stages, we performed deconvolution of bulk RNA-seq data from P10 *Jag1^Ndr/Ndr^* and *Jag1^+/+^* livers (Andersson *et al*, 2018), using a neo- and post-natal liver scRNA-seq reference (Liang *et al*, 2022). Neonatal hepatocytes were significantly depleted in *Jag1^Ndr/Ndr^* livers (**Fig. 3E**). We then estimated a maturation stage of the hepatocytes in P10 livers using deconvolution with liver time-course scRNAseq datasets ((Liang *et al*, 2022), **Fig 3F**) and ((Yang *et al*, 2023), **Fig. 3G**). Both analyses showed that *Jag1^+/+^* P10 livers are, as expected, enriched for stage-matched hepatocytes (P7 for Liang et al., 2022 and P9 – P15 for Yang et al., 2023.

**Figure 3:**
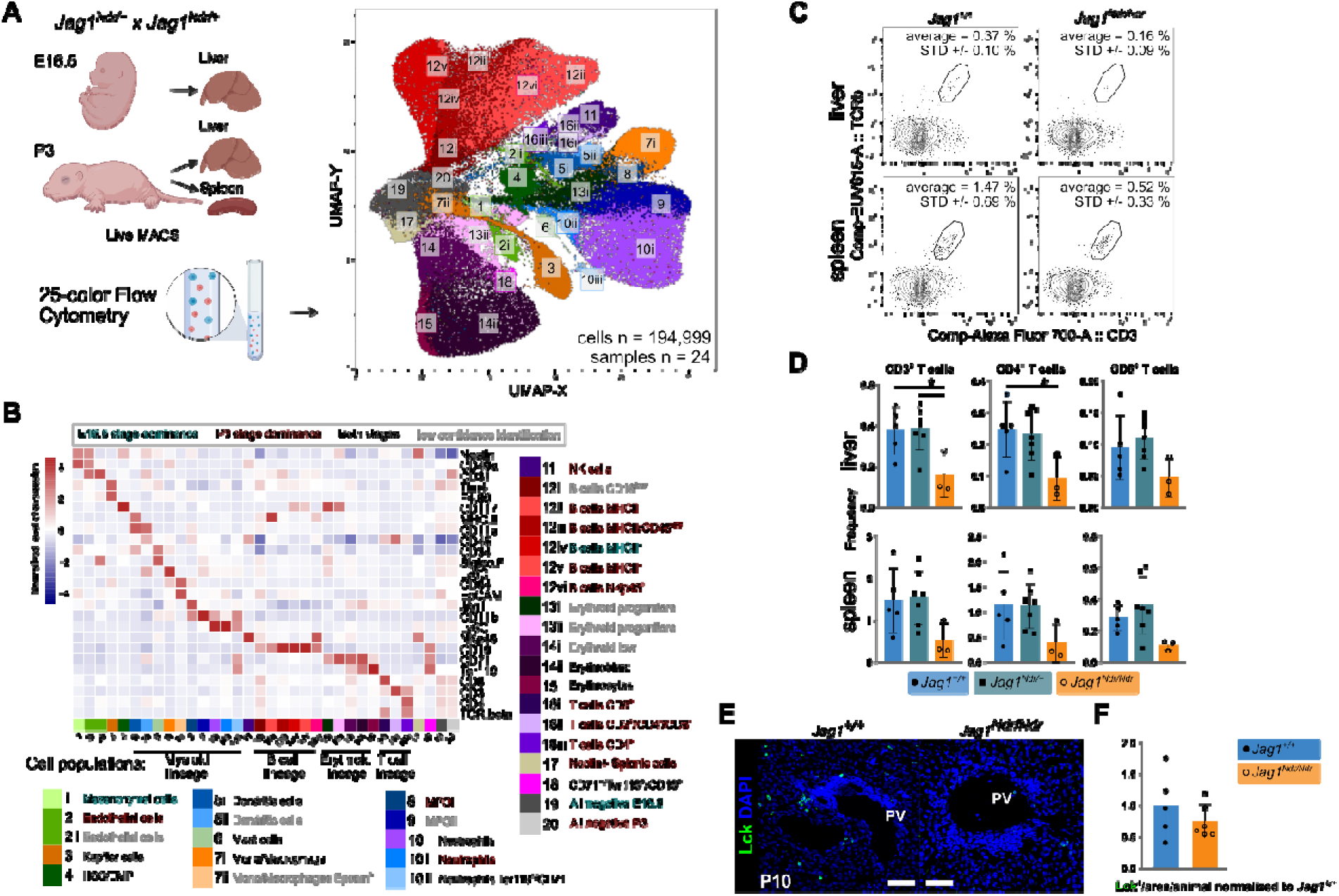
Flow cytometry analyses reveal a reduction of T cells in Jag1^Ndr/Ndr^ liver. (**A**) UMAP projection of randomly selected 194,999 cells analyzed by 25-color flow cytometry, with cell type-annotated clusters, matched to (Fig. 2A) insofar as possible. Livers were collected at E16.5 or P3, and spleens were collected at P3. (**B**) Heatmap showing a column z-score of median protein expression levels of cell type markers in the aggregated 25-colour flow cytometry dataset from E16.5 and P3 livers, and P3 spleens of *Jag1^+/+^*(n=7), *Jag1^Ndr/+^*(n=11), and *Jag1^Ndr/Ndr^* (n=6) animals. (**C,D**) Representative flow cytometry plots (C) and relative frequency of the CD3^+^, CD4^+^/CD3^+^, and CD8^+^/CD3^+^ T cells in livers and spleens from the *Jag1^+/+^*, *Jag1^+/Ndr^* and *Jag1^Ndr/Ndr^* mice at P3 (D). (**E,F**) Representative immunohistochemical staining (E) and quantification (F) of LCK^+^ T-cells in periportal areas in *Jag1^Ndr/Ndr^* and *Jag1^+/+^* livers at P10. PV, portal vein. All graphs represent mean ± SD. Statistical analysis in (C) was performed by unpaired, two-tailed Student’s t-test, *, p <0.01); Analysis in (E) is one-way ANOVA with Dunnett’s multiple comparison test, p ≤ 0.05 = *. Each data point represents a biological replicate.

In contrast, Jag*1^Ndr/Ndr^* P10 livers were enriched for less mature hepatocytes, with 68% ^+/-^6% constituted of P3 hepatocytes using Liang et al., 2022. Deconvolution using Yang et al., 2023, which includes a wider range of timepoints (E17.5 – P60), revealed a variability in proportions of different maturation stages between Jag*1^Ndr/Ndr^* P10 liver samples. Interestingly signatures of both E17.5, and P30 hepatocytes were represented in the data, possibly indicating an injury response rather than differentiation defect.

An “Immature hepatocyte” phenotype arises in adult hepatocytes upon injury (Ben-Moshe *et al*, 2022; Iwai *et al*, 2000; Nakano *et al*, 2017). To identify genes driving the immature hepatocyte signature of Jag*1^Ndr/Ndr^* P10 livers, we generated a reference gene signature for hepatoblasts and hepatocytes using a published bulk RNA-seq dataset of isolated primary murine hepatoblasts and hepatocytes (top 500 enriched genes, Supplementary table **ST6,7** from (Belicova *et al*, 2021)), and performed gene set enrichment analysis (GSEA) with P10 liver bulk RNA-seq (**Fig. 3H**). *Jag1^Ndr/Ndr^* P10 livers were enriched for hepatoblast signature genes (**Fig. 3H** left**, I**,) and depleted for hepatocyte signature genes (**Fig. 3H** right**, I**). Furthermore, the hepatoblast marker alpha fetoprotein (AFP) was 3.1-fold enriched (**Fig. 3J,K**), while the mature hepatocyte marker CYP1A2 protein was 1.7-fold less expressed (**Fig. 3L-M**). The *Cyp1a2* and additional maturation marker *Glu1* decrease was also present at the level of mRNA (**Fig. 3N**). Finally, we analyzed bulk RNA-seq data of livers from people with ALGS and found a similar enrichment of hepatoblast signature genes and down-regulation of hepatocyte-related genes in ALGS (**Fig. 3H,I**), suggesting hepatocytes are immature in both *Jag1^Ndr/Ndr^* mice and in ALGS.

To see if the immature hepatocytes signature points at hepatocyte injury response, we tested whether the immature *Jag1^Ndr/Ndr^* hepatocytes become activated and initiate a pro-inflammatory Egr1-driven response upon cholestatic injury (Allen *et al*, 2011). Re-analysis of the P10 bulk RNAseq dataset showed that pro-inflammatory *Egr1* was significantly but only mildly upregulated in *Jag1^Ndr/Ndr^* mice (2-fold), as were its targets *Cxcl1* (5.1-fold), *Cxcl2* (4-fold), *Cxcl10* (3.5-fold), *Ccl2* (4.7-fold), *Serpine1* (6.3-fold), *Isg20* (2.3-fold), and *Plaur* (2.9-fold). GSEA revealed a significant enrichment of myeloid leukocyte signatures in *Jag1^Ndr/Ndr^* liver but only modest lymphocytic contribution, indicating the mild hepatocyte activation may fail to attract lymphocytes (**Fig. 3O, ST8**). In sum, scRNA seq, bulk RNA seq, and histological analyses revealed that hepatocytes are immature in *Jag1^Ndr/Ndr^* mice and in people with ALGS. The mild pro-inflammatory activation of immature hepatocytes could underlie the inability of *Jag1^Ndr/Ndr^* liver to attract and/or activate T cells upon cholestatic injury.

### Intrahepatic T cells are underrepresented in Jag1^Ndr/Ndr^ mice

The injured liver attracts immune cells to repair tissue damage, clear dead cells, and restore homeostasis (Hammerich & Tacke, 2023) and hepatocyte activation is crucial in this process (Allen *et al*, 2011). To investigate the immune cell composition of *Jag1^Ndr/Ndr^* mice, we analyzed livers and spleens with 25-color flow cytometry (**Fig. 3A**). Unsupervised clustering identified 20 cell types in 34 sub-populations, including mesenchymal cells, endothelial cells, and a majority of hematopoietic cells (**Fig. 3A,B, S3A-C**). There was a tendency towards fewer T cells in *Jag1^Ndr/Ndr^* spleens at P3, and a significant 61% decrease in the frequency of CD4^+^ T cells in *Jag1^Ndr/Ndr^*cholestatic livers compared to healthy *Jag1^+/+^* controls of the same stage (**Fig 3C,D, S3D**). We next stained and quantified lymphocyte-specific protein tyrosine kinase positive (LCK^+^) T cells in periportal areas of *Jag1^Ndr/Ndr^*livers at P10, at which point *Jag1^Ndr/Ndr^* mice are highly cholestatic (Andersson *et al*, 2018). Despite the ongoing cholestasis that should attract immune cells (Kisseleva & Brenner, 2021), there was no enrichment of LCK^+^ T cells in *Jag1^Ndr/Ndr^* portal areas (**Fig. 3E,F**). The liver and splenic immune cell composition in *Jag1^Ndr/Ndr^* mice thus suggest a defect in T cell response to cholestasis which could impact the fibrotic process.

### Disrupted thymic development and altered T cell differentiation in Jag1^Ndr/Ndr^ mice

Our data suggests that hepatocytes fail to attract T cells to the liver, but the T cell development could also be disrupted in *Jag1^Ndr/Ndr^* mice (**Fig. 3D-F**). Notch signaling modulates immune system development (Vanderbeck & Maillard, 2021), and the immune system modulates the fibrotic process (Kisseleva & Brenner, 2021), but it is not known whether the immune system is impacted in ALGS and if it can affect fibrosis. We therefore next investigated the immune system in *Jag1^Ndr/Ndr^*mice, its competence, and its role in liver fibrosis.

First, we characterized hematopoiesis and T cell development in *Jag1^Ndr/Ndr^*mice at embryonic and postnatal stages. Hematopoietic progenitor development is regulated by Jag1-Notch signaling (Robert-Moreno *et al*, 2008). However, flow cytometry of E9.5 *Jag1^Ndr/Ndr^* yolk sac and embryo proper revealed no differences in erythroid, macrophage, megakaryocyte, or erythro-myeloid progenitor populations at this stage (**Fig. S4A-D**), suggesting the first wave of definitive immune cell production is unaffected in *Jag1^Ndr/Ndr^* mice.

Lymphocyte progenitors differentiate into T cells in the thymus, a primary lymphoid organ in which thymic epithelial cells (TECs) provide instructive cues (Brandstadter & Maillard, 2019). Thymocytes that are double negative (DN) for CD4 and CD8 develop into double positive thymocytes (DP) which subsequently differentiate into CD8+ or CD4+ T cells (Brandstadter & Maillard, 2019). The CD4+ T cells can further develop into regulatory CD4+ T cells (Tregs) (**Fig. 4A**). Several of these steps are Notch-dependent (Brandstadter & Maillard, 2019), but whether they are affected in ALGS is not known. We therefore assessed the thymus and its constituent cell types in *Jag1^Ndr/Ndr^* mice.

**Figure 4:**
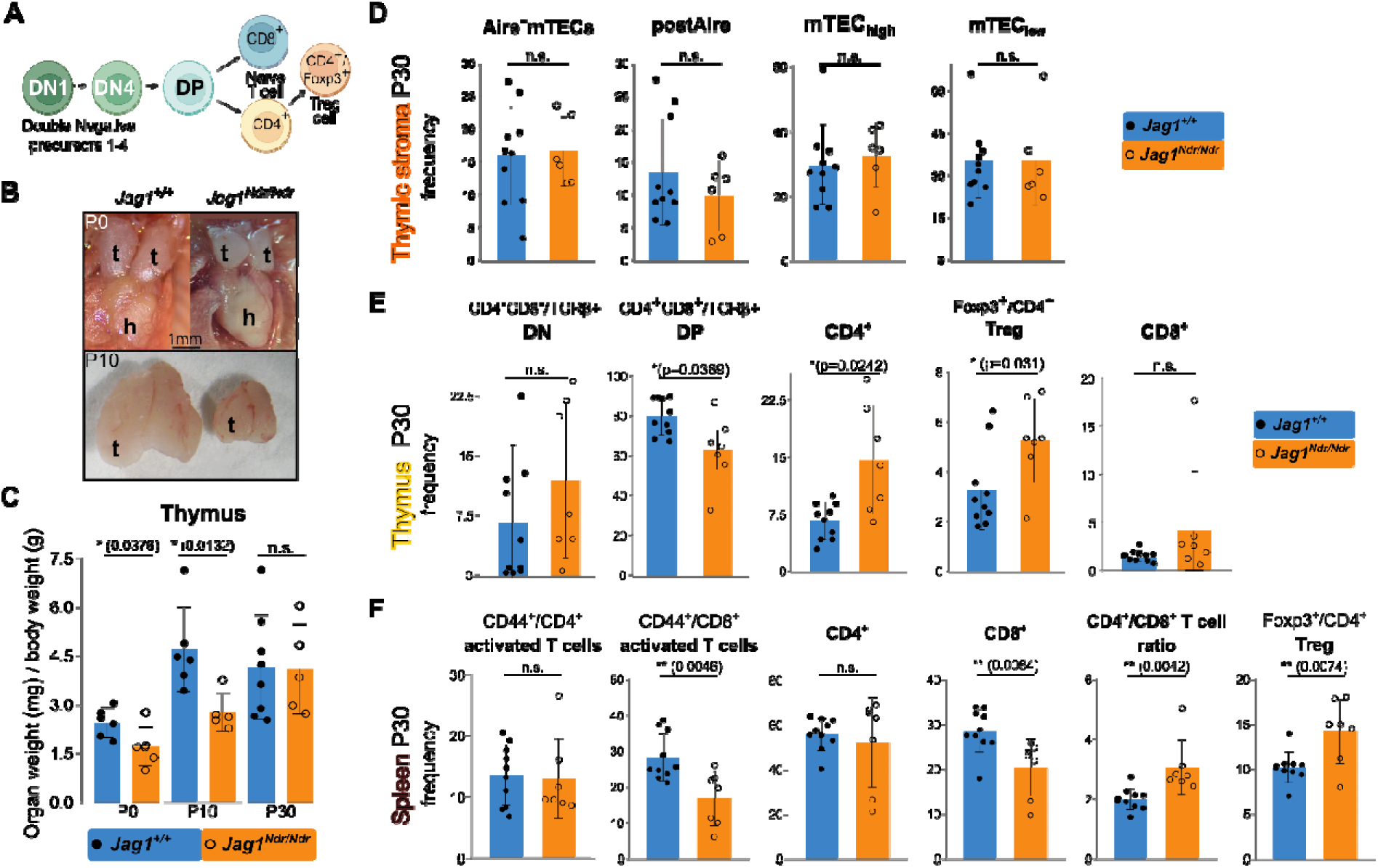
Thymic developmental delay and excess Tregs in postnatal Jag1^Ndr/Ndr^ mice. (**A**) Schematic of thymocyte development in the thymus. (**B**) Macroscopic images of *Jag1^+/+^* and *Jag1^Ndr/Ndr^* thymi at P0 and P10. (**C**) Thymic weights at P0, P10 and P30, normalized to body weight. (**D-F**) Flow cytometric analyses of cell frequencies of thymic epithelial cells, TECs (D), thymic thymocytes/T cells (E), and splenic T cells (F) in *Jag1^+/+^*and *Jag1^Ndr/Ndr^* mice at P30. Each data point represents a biological replicate. Mean ± SD, Unpaired, two-tailed Student’s t-test, p ≤ 0.05 = *, p ≤ 0.01 = **, n.s. not significant. t, thymus; h, heart; DN, Double Negative; DP, Double Positive; Mac, macrophage; NK, Natural Killer cell; TEC, Thymic Epithelial Cell; mTEC, medullary TEC; cTEC, cortical TEC;Treg, regulatory T cell

*Jag1^Ndr/Ndr^* thymi were significantly smaller than *Jag1^+/+^* thymi at both P0 and P10 but achieved a normal size by P30 (**Fig. 4B,C**). We mapped the expression of Notch components in human and mouse developing thymus using a published scRNA-seq atlas encompassing developmental stages from fetal, pediatric, and adult human individuals, and mouse thymi at 4, 8, and 24 weeks of age (Park *et al*, 2020). This analysis confirmed enriched expression of *Jag1* in medullary Thymic Epithelial Cells (mTECs) (Li *et al*, 2020), and Notch1-mediated activation (high *Notch1*/*Hes1* expression) in DN thymocytes prior to differentiation into CD4^+^ or CD8^+^ T cells (Lehar *et al*, 2005; Radtke *et al*, 2004) (**Fig. S4E,F**). However, although *Jag1* is expressed in TEC subpopulations and Notch signaling is active in these cell types with upregulated *Hes1* and *Nrarp* expression (**Fig. S4E,F**), there were no significant differences in *Jag1^Ndr/Ndr^* TEC subpopulations including Aire^+^ mTECs, postAire mTECs (MHCII^+^Ly6D^+^Aire^neg/low^), mTECs^low^ (MHCII^low^CD80^low^) or mTECs^high^ (MHCII^high^CD80^high^) in *Jag1^Ndr/Ndr^* mice (**Fig. 4D**), suggesting that while *Notch1* regulates TEC specification (Li *et al*, 2020), Jag1^Ndr^ does not affect this process.

We next analyzed T cell differentiation in the thymus at P30. While the frequency of DN thymocytes was similar in *Jag1^+/+^* and *Jag1^Ndr/Ndr^* thymi, we detected ¼ reduction of DP thymocytes frequency in *Jag1^Ndr/Ndr^* thymi, suggesting skewed differentiation in *Jag1^Ndr/Ndr^* mice (**Fig 4E**). In contrast, CD4^+^ T cells and Tregs were twice as prevalent in *Jag1^Ndr/Ndr^* thymi, while the CD8^+^ T cell frequency was unaffected in the *Jag1^Ndr/Ndr^* thymus (**Fig. 4E**). These data suggest a disrupted DN-to-DP differentiation, and excess production of T cells, particularly Tregs.

Finally, we assessed immune composition in the spleen, to determine how defects established in the thymus continue to develop in this secondary lymphoid organ and investigate whether *Jag1^Ndr/Ndr^* immune cells are correctly activated. While there were no differences in CD4^+^ or CD44^+^/CD4^+^ activated T cell populations in *Jag1^Ndr/Ndr^* spleens, there were significantly fewer CD8^+^ T cells and CD44^+^/CD8^+^ activated T cells (**Fig. 4F**). The decrease in CD8+ T cells led to an increased CD4^+^/CD8^+^ T cell ratio in *Jag1^Ndr/Ndr^* spleens. Importantly, similar to the thymus, Tregs were 39% more frequent in the *Jag1^Ndr/Ndr^* spleens (**Fig. 4F**). Collectively, these data revealed a role for Jag1 in thymic development in a model for ALGS, apparent already at P0, preceding cholestasis. We therefore next aimed to investigate the functional competence of the *Jag1^Ndr/Ndr^* immune lymphoid compartment.

### Jag1^Ndr/Ndr^ T cells are less inflammatory

To investigate the function of *Jag1^Ndr/Ndr^* T cells, we first performed lymphocyte adoptive transfer experiments into *Rag1^-/-^* mice, which lack mature B- and T cell populations, but provide a host environment with normal Jag1 (Mombaerts *et al*, 1992). T cell transfer to lymphodeficient recipients leads to mild gastrointestinal tract inflammation, the degree of which reflects hyper- or hypo-active T cells in homeostatic conditions. *Rag1^-/-^* mice were transferred with *Jag1^Ndr/Ndr^* or *Jag1^+/+^* lymphocytes, their weight was monitored, and after 8 weeks the intestine was analyzed histologically, and the mesenteric gland was analyzed with flow cytometry (**Fig. S5A**). There were no significant differences in survival, weight, or intestinal inflammatory status, between the recipients of the *Jag1^Ndr/Ndr^*and *Jag1^+/+^* lymphocytes (**Fig. S5B-E**), demonstrating that *Jag1^Ndr/Ndr^* lymphocytes are neither hyper- nor hypo-active in homeostatic conditions.

We next assessed *Jag1^Ndr/Ndr^* T cell capacity to respond to challenge, using the well-established recovery model of Dextran Sodium Sulphate (DSS)-induced intestinal colitis (Chassaing *et al*, 2014). Four weeks after *Jag1^+/+^*→*Rag1^-/-^* or *Jag1^Ndr/Ndr^*→*Rag1^-/-^*lymphocyte transfer, mice were challenged with two consecutive 10 day-long 2.5% DSS diets, inducing bacterial infiltration and inflammation of the intestinal submucosa (Hernández-Chirlaque *et al*, 2016), followed by 7 days of tissue recovery, after which the intestine was analyzed histologically, and the mesenteric lymph node by flow cytometry (**Fig. 5A**). The intestines and colons were shorter and heavier in *Jag1^Ndr/Ndr^* lymphocyte recipients, demonstrating less efficient recovery (**Fig. 5B**). Histological scoring of intestinal and colonic inflammation revealed significantly less inflammation in *Jag1^Ndr/Ndr^ Rag1^-/-^* animals (**Fig. 5C,D**). Finally, *Jag1^Ndr/Ndr^* lymphocyte transfer resulted in significantly fewer activated CD44^+^/CD4 ^+^ and CD44^+^/CD8^+^ T cells in the mesenteric lymph node compared to *Jag1^+/+^*→ *Rag1^-/-^* controls (**Fig. 5E**). In sum, these adoptive transfer experiments show that *Jag1^Ndr/Ndr^* T cells behave normally in homeostatic conditions but are less activated and less efficient at mediating recovery from intestinal insult. The *Jag1^Ndr/Ndr^* lymphocyte altered ability to become activated or sustain inflammation could impact liver fibrosis, which we next investigated.

**Figure 5:**
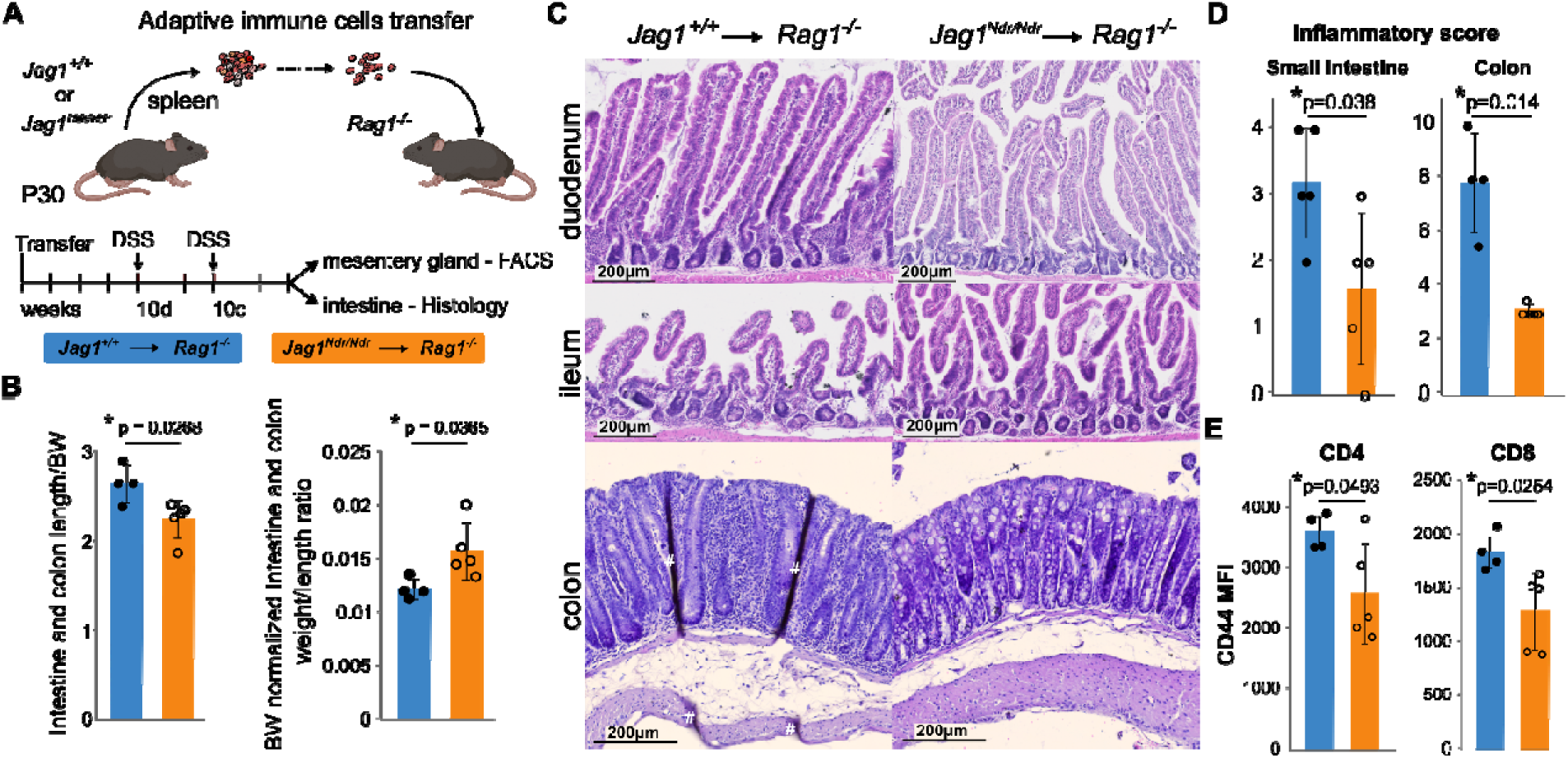
Jag1^Ndr/Ndr^ T cells do not mount an adequate response to DSS-induced colitis. (**A**) Scheme of the Dextran sodium sulfate (DSS)-induced colitis. (**B**) Length, and weight/length ratios of the intestines and colons from DSS-treated *Jag1^+/+^*→*Rag1^-/-^* and *Jag1^Ndr/Ndr^*→*Rag1^-/-^* mice, normalized to body weight. (**C,D**) Representative H&E-stained intestinal (top), and colonic (bottom) sections from the *Jag1^+/+^* or *Jag1^Ndr/Ndr^* T cell-transferred *Rag1^-/-^* mice after DSS treatment (C), and their inflammatory scores (D). (**E**) Flow cytometry analysis of the mean fluorescence intensity (MFI) of CD44 staining of CD4^+^ and CD8^+^ T cells from DSS-treated *Jag1^+/+^*→*Rag1^-/-^* and *Jag1^Ndr/Ndr^*→*Rag1^-/-^*mesenteric lymph nodes. (Each data point represents a biological replicate. mean ± SD, Unpaired, two-tailed Student’s t-test, p ≤ 0.05 = *). # - tissue-folding artefact

### Jag1^Ndr/Ndr^ T cells attenuate BDL-induced cholestatic fibrosis

Tregs are anti-fibrotic in the context of the cholestatic liver injury (Roh *et al*, 2014; Zhang & Zhang, 2020), and were enriched in *Jag1^Ndr/Ndr^*mice (**Fig. 4**), which could attenuate the CD4^+^ and CD8^+^ response to intestinal insult (**Fig. 5**). W therefore next tested *Jag1^Ndr/Ndr^*T -cell effects on liver fibrosis. We investigated *Jag1^Ndr/Ndr^*immune cell contribution to fibrosis using bile duct ligation (BDL), a surgically induced obstructive cholestasis model involving neutrophils and T cells (Licata *et al*, 2013). The transplanted *Rag1^-/-^* recipients were subjected to BDL surgery four weeks after *Jag1^+/+^*or *Jag1^Ndr/Ndr^* lymphocyte transfer (**Fig. 6A**). BDL-treatment increased total and conjugated bilirubin, alkaline phosphatase (ALP), aspartate transaminase (AST) and alanine transaminase (ALT), reflecting ongoing liver damage, irrespective of which lymphocytes were transplanted (physiological levels in grey, **Fig. 6B)**.

**Figure 6:**
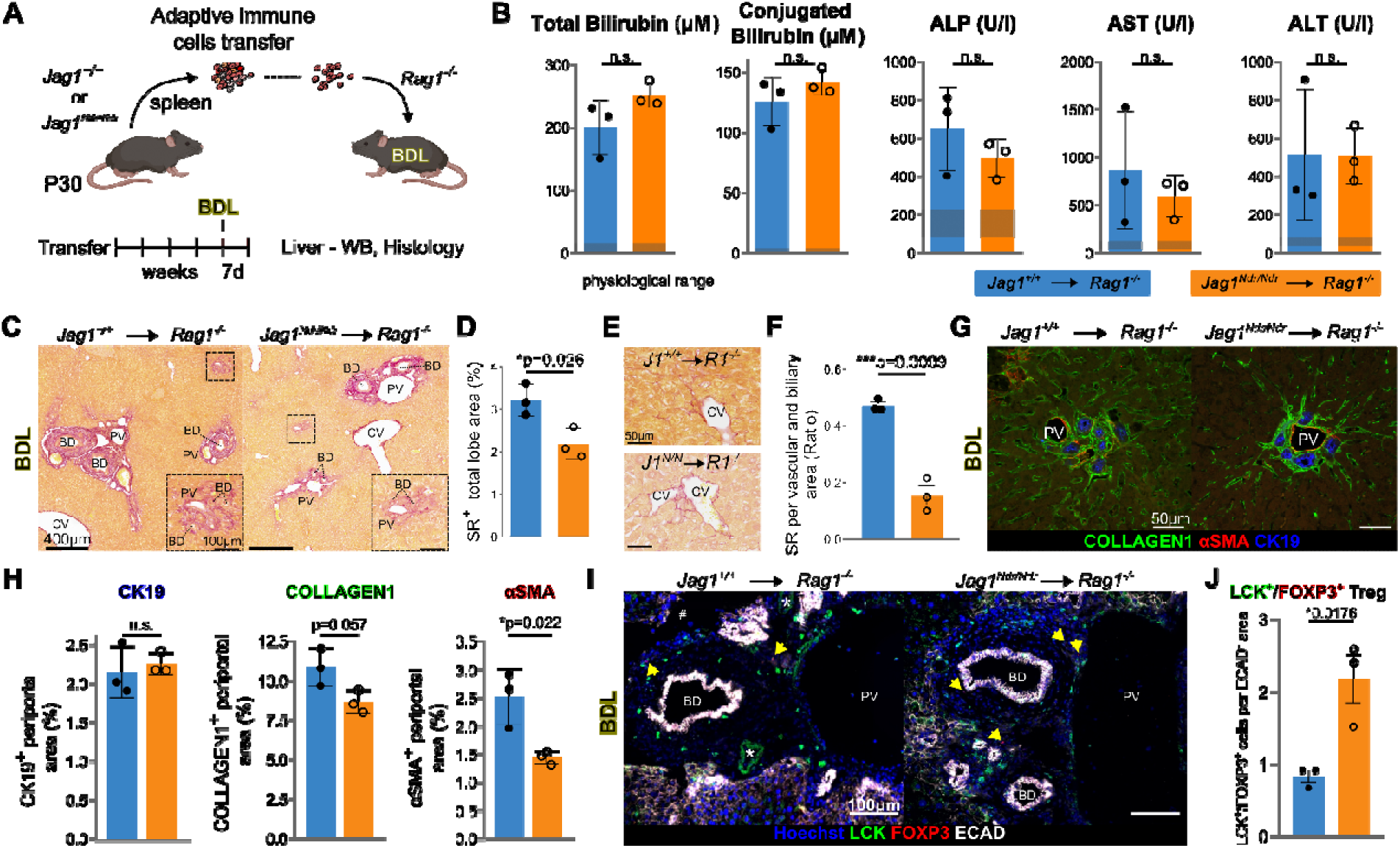
*Jag1^Ndr/Ndr^* ^T^regs limit the extent of BDL-induced periportal fibrosis. (**A**) Scheme of the BDL experimental model. (**B**) Liver biochemistry of the *Jag1^+/+^* →*Rag1^-/-^* and *Jag1^Ndr/Ndr^* →*Rag1^-/-^* mice after BDL. Grey area indicates physiological ranges. (**C,D**) Representative images of sirius red (SR) staining of *Jag1^+/+^* →*Rag1^-/-^* (left) and *Jag1^Ndr/Ndr^* →*Rag1^-/-^* (right) liver BDL (C), and corresponding quantification (D). (**E**) Representative SR staining in Zone 3 of *Jag1^+/+^* →*Rag1^-/-^* and *Jag1^Ndr/Ndr^* ^→^*Rag1^-/-^* mice after BDL. (**F**) Quantification of SR staining in periportal areas of *Jag1^+/+^* →*Rag1^-/-^* and *Jag1^Ndr/Ndr^* →*Rag1^-/-^* livers after BDL. (**G,H**) CK19, SMA, and Collagen1 staining of *Jag1^+/+^* →*Rag1^-/-^* and *Jag1^Ndr/Ndr^* →*Rag1^-/-^* livers after BDL (G), and quantification in periportal areas (H). (**I,J** Representative immunofluorescent images of liver from *Jag1^+/+^*→*Rag1^-/-^* (left) and *Jag1^Ndr/Ndr^*→*Rag1^-/-^* (right) mice after BDL, stained with α-LCK and α-FOXP3, ECAD and Hoechst, yellow arrows mark LCK^+^/FOXP3^+^ Tregs (I), with LCK^+^/FOXP3^+^ Treg quantification in ECAD^-^ periportal area (J). (mean ± SD, unpaired, two-tailed Student’s t-test); # - tissue-folding artefact; CV, central vein; PV, portal vein; BD, bile duct; HA, hepatic artery.

To test immune cell impact on the extent of fibrosis, we analyzed collagen deposition with SR staining. Fibrosis levels in BDL-treated *Jag1^Ndr/Ndr^*→*Rag1^-/-^*livers were significantly lower than in BDL-treated *Jag1^+/+^*→*Rag1^-/-^* livers (**Fig. 6C,D**). Importantly, the impact on fibrosis was region-specific: pericentral fibrosis, with pericellular and perisinusoidal fibrosis, was similar in *Jag1^+/+^*→*Rag1^-/-^*and *Jag1^Ndr/Ndr^*→*Rag1^-/-^* mice (**Fig. 6E, S6A**), while periportal fibrosis was reduced by threefold in *Jag1^Ndr/Ndr^*→*Rag1^-/-^* livers (**Fig. 6F**). To determine whether *Jag1^Ndr/Ndr^* lymphocytes affect fibrotic response to BDL, we analyzed periportal expression of cytokeratin 19 (CK19, a marker of cholangiocytes), alpha smooth muscle *al*, 2014)). The CK19^+^ ductular reaction area was similar in *Jag1^+/+^*→*Rag1^-/-^*and *Jag1^Ndr/Ndr^*→*Rag1^-/-^* mice subjected to BDL. However, there was a tendency towards less collagen1, and there was significantly less aSMA^+^ staining in *Jag1^Ndr/Ndr^*→*Rag1^-/-^* livers subjected to BDL (**Fig. 6G,H**).

Next, we assessed T cell liver infiltration upon BDL-induced cholestatic liver injury with western blot and immunostaining for LCK. There were no differences in the capacity of *Jag1^Ndr/Ndr^* or *Jag1^+/+^* T cells to migrate into the liver upon BDL-induced liver injury (**Fig. S6B,C**), with numerous LCK^+^ T cells present in BDL ductular reaction loci (**Fig. S6D,E**). Intriguingly, there was a 2-fold increase in LCK^+^/FOXP3^+^ double positive Treg cells in the periportal area of *Jag1^Ndr/Ndr^*→*Rag1^-/-^* livers subjected to BDL (**Fig. 6I,J**).

To determine whether there could be a relationship between immune cell composition and fibrosis in patients with ALGS, we again analyzed previously published bulk RNAseq data (Andersson *et al*, 2018). A biliary stricture gene signature, reflecting cholestasis, was mildly upregulated in ALGS samples, while both immature T cell and Treg (FOXP3-driven) gene signatures were significantly enriched in livers from patients with ALGS (**Fig. S6F**). Interestingly, *COL1A1* expression was tightly and positively correlated with *FOXP3* levels in non-ALGS liver disease (R^2^=0.9904), suggesting that increased fibrosis (*COL1A1*) is associated with increased Treg presence (*FOXP3* expression) in other liver diseases. However, in ALGS, *COL1A1* was inversely correlated with *FOXP3* levels, suggesting that an increase in Tregs may be associated with less extensive fibrosis in ALGS (**Fig. S6G)**. These data corroborate that intrahepatic lymphocytes could be affected in ALGS and that their presence could modify the course of fibrosis in an ALGS-specific manner.

In conclusion, results from BDL experiments demonstrate that both *Jag1^+/+^* and *Jag1^Ndr/Ndr^* lymphocytes can home to a cholestatic liver, but that *Jag1^Ndr/Ndr^* and possibly also ALGS Treg cells modify the periportal fibrotic response.

## DISCUSSION

Inflammation and fibrosis impair liver function, harm hepatocytes, and can initiate a cascade of events leading to portal hypertension, and ultimately liver cancer (Henderson *et al*, 2020). Notch signaling regulates liver development, and its disruption results in ALGS and bile duct paucity (Kohut *et al*, 2021). Whether Notch disruption in ALGS affects immune system development, and thus alters the course of fibrotic liver disease, was not known. Here, we dissected the onset of fibrosis using unbiased single-cell techniques, and transcriptomic cross-comparisons between *Jag1^Ndr/Ndr^* and ALGS patient datasets. We show that cholestatic *Jag1^Ndr/Ndr^* mice recapitulate ALGS-like pericellular fibrosis (Fabris *et al*, 2007), and develop splenomegaly, a common consequence of portal hypertension. Our unbiased omics analysis of the neonatal cholestatic liver revealed that *Jag1^Ndr/Ndr^* hepatocytes exhibit an immature phenotype, with a limited capacity to transform into the activated pro-inflammatory state. Further, we showed that T cell development and function is affected in *Jag1^Ndr/Ndr^* mice. We tested the competence of *Jag1^Ndr/Ndr^*lymphocytes in a series of transplantation experiments, demonstrating that *Jag1^Ndr/Ndr^*lymphocytes’ activation and contribution to periportal but not parenchymal fibrosis is limited, likely via aberrant activity of Tregs. Jag1 thus regulates hepatocyte injury response and thymocyte development and competence, with implications for the course of liver disease in ALGS.

*Jag1^Ndr^* mice are a suitable model to explore the consequences of systemic Jag1 dysregulation, which is relevant in ALGS (Andersson *et al*, 2018; Hankeova *et al*, 2021, 2022; Iqbal *et al*, 2024; Mašek & Andersson, 2024). As such, the immature hepatocytes in *Jag1^Ndr/Ndr^* mice could either arise due to differentiation defects, or in response to liver injury. Adult hepatocytes revert to an “immature hepatocyte phenotype” upon injury (Ben-Moshe *et al*, 2022; Iwai *et al*, 2000; Nakano *et al*, 2017), and in the context of CCL4 injury, JAG1 expressed by activated HSCs induces NOTCH2-dependent dedifferentiation of mature hepatocytes into Afp-expressing immature hepatocytes (Nakano *et al*, 2017). Whether an immature hepatocyte phenotype can be triggered by injury even in a Jag1 insufficiency context is thus an intriguing question. While the JAG1^NDR^ protein cannot bind or activate NOTCH1 (Hansson *et al*, 2010), it retains some capacity to bind and activate NOTCH2 (Andersson *et al*, 2018), which may be sufficient to induce dedifferentiation upon injury. It would be of interest to test whether Jag1 potency affects injury-induced reversion of hepatocytes in patients as a function of their genotype.

Despite acute cholestasis at P3 and P10, there was only modest or no induction of distress molecules or chemokines that are otherwise upregulated in activated hepatocytes (Allen *et al*, 2011). *Egr1* is an early stress response gene (Gashler & Sukhatme, 1995), and its absence strongly attenuates inflammation in the BDL model of cholestasis (Allen *et al*, 2011). The reduced expression of *Egr1* in E16.5 *Jag1^Ndr/Ndr^* hepatocytes is in line with its decrease in hematopoietic stem cells in *Pofut1* KO mice. In this model the reduction in *Egr1* is attributed not to Notch activation status but to impaired adhesion of Notch receptor-expressing hematopoietic stem cells to ligand-expressing neighboring cells, normally mediated by fucose-dependent Notch-receptor-ligand engagement (Wang *et al*, 2015). *Egr1* may thus be Jag1/Notch-regulated, limiting the capacity of hepatocytes to become activated in the *Jag1^Ndr/Ndr^* environment with reduced Jag1 activity (**Fig. ST2, 2O**).

While fibrosis is the focus of this work, the reduced CD4^+^CD8^+^ DP cells and the CD4^+^ Treg enrichment in *Jag1^Ndr/Ndr^* mice is intriguing given the prominent roles of Notch in immune cell development (Brandstadter & Maillard, 2019). A systematic assessment of the immune system in patients with ALGS in a large, controlled cohort of patients has not been reported, although current data suggest the immune system may be compromised, with frequent (re)infections in ∼25% of patients (Tilib Shamoun *et al*, 2015). *In vitro* co-culture experiments revealed that low Notch activation by JAG1 exhibits stage-specific functions in DN thymocyte development and induces CD4^+^ T cell differentiation rather than DP differentiation (Lehar *et al*, 2005). *In vivo*, *Jag1* overexpression in T-lymphocyte progenitors (Beverly *et al*, 2006) or conditional *RBPj* silencing in TECs (García-León *et al*, 2022) induces TEC apoptosis and drives premature thymic involution. In *Jag1^Ndr/Ndr^* mice a small-for-size thymus was evident at P0 and P10 (**Fig. 4B,C**) which normalized by P30.

Whether the thymus size normalization at P30 is due to inhibited TEC apoptosis (**Fig. 4C**), and whether this impacts thymic involution at later stages, is an interesting question that warrants further investigation. Nonetheless, the small *Jag1^Ndr/Ndr^* thymus and hypomorphic TEC-mediated JAG1 signaling (Lehar *et al*, 2005) could explain the dearth of CD4+ T cells at P3 in *Jag1^Ndr/Ndr^* livers (**Fig 3C-F**). Interestingly, JAG1 can induce differentiation of T cells into Tregs (Vigouroux *et al*, 2003; Hoyne *et al*, 2000), which implies Tregs should be reduced in *Jag1^Ndr/Ndr^* mice. Instead, Tregs were more frequent in both *Jag1^Ndr/Ndr^* thymus and spleen at P30 (**Fig. 4E,F**). The enrichment in Tregs in *Jag1^Ndr/Ndr^* mice is, however, in line with the reported Treg expansion upon *Notch1* or *RBPj* inactivation (Charbonnier *et al*, 2015). Finally, a Notch-independent Jag1-CD46 interaction was proposed to instruct Th1 responses in T cells (Le Friec *et al*, 2012), a process that was inhibited in patients with ALGS (Le Friec *et al*, 2012), but it is unknown if any analogous mechanism could also impact T cell development. Further studies to conditionally abrogate *Jag1 in vivo* would help further delineate the temporal and spatial requirements for *Jag1* in Treg development in ALGS.

The transplantation of *Jag1^Ndr/Ndr^* immune cells resulted in reduced periportal fibrosis compared to transplantation of *Jag1^+/+^*immune cells (**Fig. 6**). In this experiment, the recipient *Rag1^-/-^* mice have normal hepatic Notch signaling, and only Notch-defective immune cells are transplanted. The comparable numbers of LCK^+^ T cells in the livers of transplanted *Rag1^-/-^* mice (**Fig. S6B-E**) demonstrate that *Jag1^Ndr/Ndr^* T cells can respond to chemoattraction by activated hepatocytes, corroborating our interpretation that the *Jag1^Ndr/Ndr^* hepatocytes are unable to adequately attract lymphocytes due to their low/absent inflammatory signature (**Fig. 2O**). The *Jag1^Ndr/Ndr^* immune cells present three T cell phenotypes which all could mitigate liver fibrosis: fewer and less activated CD8^+^ T cells which would otherwise promote liver fibrosis (Shivakumar *et al*, 2007) (**Fig 4F**) and an increase in Tregs, which restrict liver fibrosis (Zhang & Zhang, 2020; Roh *et al*, 2014) (**Fig 4E, F**). Importantly, while hepatic stellate cells are major drivers of fibrosis, portal fibroblasts appear to play a relatively stronger role in BDL-induced fibrosis (Iwaisako *et al*, 2014; Lua *et al*, 2016; Koyama *et al*, 2017). The fact that *Jag1^Ndr/Ndr^* lymphocytes limit periportal fibrosis after the BDL suggests that portal fibroblast-induced fibrosis may be limited by *Jag1*-deficient T cell populations. Resolving whether HSCs and/or portal fibroblasts are key drivers of fibrosis in ALGS would be important for further studies on the interaction between the immune system and myofibroblasts in ALGS.

The pericellular fibrosis in *Jag1^Ndr/Ndr^* mice recapitulates the fibrotic phenotype of patients with ALGS and suggests that *Jag1* is required for mounting a strong, or rapid, periportal fibrotic response, as seen in biliary atresia (Fabris *et al*, 2007). Fibrotic liver repair in ALGS is characterized by an expansion of hepatobiliary cells but no overt increase in reactive ductular cells, while the highly fibrotic biliary atresia is characterized by an increase in reactive ductular and hepatic progenitor cells (Fabris *et al*, 2007). Reactive ductular cells are pro-fibrotic (Banales *et al*, 2019), and their absence in ALGS may thus limit bridging or periportal fibrosis (Fabris *et al*, 2007). The transplantation experiments described here show that *Jag1-*compromised lymphocytes also contribute to attenuated periportal fibrosis. However, Notch signaling modulates fibrosis via multiple other cell types in the liver, which may also play a role in ALGS. Notch activation in hepatocytes drives fibrosis in the context of NAFLD, and its inhibition attenuates NAFLD-associated fibrosis (Schwabe *et al*, 2020). Hepatocytic Jag1 also contributes directly to liver fibrosis and is required for NASH-induced liver fibrosis (Yu *et al*, 2021). Notch activation in HSCs (Bansal *et al*, 2015), or liver sinusoidal endothelial cells can also aggravate hepatic fibrosis (Duan *et al*, 2018). Intriguingly, Notch signaling is not required for methionine-choline deficient (MCD) diet-induced fibrosis (Zhu *et al*, 2018), demonstrating that fibrosis can be Notch-dependent or -independent, depending on the nature of the hepatic insult. Thus, while *Jag1^Ndr/Ndr^* immune cells are sufficient to mitigate periportal liver fibrosis in a model of biliary injury with otherwise normal Notch signaling (**Fig. 6**), multiple additional Notch-regulated mechanisms likely contribute to the fibrotic process in ALGS and remain to be investigated for their relative contribution.

In sum, we described and experimentally tested the function of the immune system in a model of ALGS. Our results demonstrate that *Jag1* mutation concurrently impacts hepatocyte maturity and adaptive immunity, and thus modulates liver fibrosis via multiple axes. We suggest that the course of liver disease in ALGS may be determined by the interaction between liver and immune system developmental defects. These results provide new insights into how Jag1 controls fibrosis in ALGS via interaction between hepatocytes and T cells.

## METHODS

### Mouse maintenance and breeding

*Jag1^Ndr/Ndr^* mice were maintained in a C3H/C57bl6 background(Andersson *et al*, 2018) and are deposited in the EMMA: https://www.infrafrontier.eu/search?keyword=EM:13207. *Rag1^-/-^*animals were purchased from Jax (Jax: 002216) and maintained in house, in compliance with the FELASA guidance in individually ventilated cages. For details, see Supplementary Methods.

### Adaptive T cell transfer and DSS-induced colitis mouse model

Adult *Rag1^-/-^* mice were used as the acceptors of T- and B-lymphocytes isolated from *Jag1^+/+^* or *Jag1^Ndr/Ndr^* mice spleens. For colitis induction by DSS (Sigma-Aldrich, #D8906) mice were given 2.5% DSS in drinking water for one week, followed by 10 days of recovery and subsequently one more week of DSS treatment. For details, see Supplementary Methods.

### Bile duct ligation model of cholestasis

Obstructive cholestasis was induced by BDL with modifications described in Supplementary material. Mice were sacrificed after 7 (BDL) for further analyses.

### Immunohistochemical procedures

Formalin-fixed, paraffin-embedded liver sections (4μm) were stained with H&E or SR. For antibodies (**Supplementary Methods Table 1**) and further details, see Supplementary Methods.

### Liver sectioning and image acquisition

Mice >P10 were euthanized with CO_2_ or <P10 were decapitated, and processed as described(Andersson *et al*, 2018). For details of staining and image acquisition see Supplementary Methods.

### Western blotting

Mouse liver tissue was weighed and homogenized in RIPA Lysis and Extraction Buffer (Thermo Scientific, #89900) with the Protease and phosphatase inhibitor (Thermo Scientific, #A32961), using 15-25mg of tissue/200μl. For details, see Supplementary Methods.

### Sample processing and flow cytometry analysis of embryo proper and yolk sac at E9.5

Dissociation and flow cytometry protocols as well as the primary antibody mixes (**Supplementary Methods Table 2**) are in Supplementary Methods.

### Flow cytometry analysis of TECs and T cells

For details, see Supplementary Methods.

### Single cell RNA-seq and FACS: E16.5 and P3 sample preparation and staining

Detailed protocol and antibodies (**Supplementary Methods Table 3)** can be found in Supplementary Methods.

### Library preparation, sequencing and analysis of the scRNA seq using Seurat packages

Demultiplexing, quality control, raw data, and gene expression data counts were performed using CellRanger (Chromium), by the BEA and SciLife core facilities and are available online (GSE236483). For details, see Supplementary Methods.

### Deconvolution of bulk RNA-seq

Bulk RNA-seq datasets (GSE104875), (GSE104873) were analyzed (Andersson *et al*, 2018), and scRNAseq dataset GSE171993 and GSE209749 (Liang *et al*, 2022; Yang *et al*, 2023) were used as references for cell type deconvolution. For details, see Supplementary Methods.

## ACKNOWLEDGEMENTS

The data handling and initial computations were enabled by resources in projects [SNIC 2018/8-200 and SNIC 2020-16-189] provided by the Swedish National Infrastructure for Computing (SNIC) at UPPMAX, partially funded by the Swedish Research Council through grant agreement no. 2018-05973. We thank the core facility at Novum, BEA, Bioinformatics and Expression Analysis, which is supported by the board of research at the Karolinska Institute and the research committee at the Karolinska hospital. We thank Vinicna Microscopy Core facility at CUNI for an excellent microscopy support. We thank Raoul Kuiper and the FENO core facility for immunohistochemical staining of JAG1 in adult mice, and for discussions. We thank Stefaan Verhulst for fruitful discussions of HSC biology and analyses which are not included in this version of the manuscript. We thank Katarina Kováčová for excellent technical support, Sandra De Haan for assistance with collection of biological material, and Roman Hillje for his easy-to-use templates for visualization in R https://romanhaa.github.io/projects/scrnaseq_workflow/. This work was supported by the following grant agencies: Grant Agency of the Czech Republic (24-10622S), PRIMUS/21/SCI/006 project funded by Charles University Grant Agency, MSCA Fellowships CZ - Charles University CZ.02.01.01/00/22_010/0002902, Wenner-Gren Foundation to JM, The European Association for the Study of the Liver (EASL: The Daniel Alagille Award to ERA and the Sheila Sherlock Post Doc fellowship to JM), funding from Karolinska Institutet (2-560/2015-280, 2-2110/2019-7, 2-195/2021), KI/SLL Center for Innovative Medicine (CIMED, 2-538/2014-29), Alex & Eva Wallström Foundation to E.R.A.,, Grant Agency of the Czech Republic (21-21736S), and ID Project No. LX22NPO5102 - Funded by the European Union - Next Generation EU to M.G. Cancerfonden fellowship for I.F. DVO, AF, TB and JD acknowledge support from Talking microbes - understanding microbial interactions within One Health framework (CZ.02.01.01/00/22_008/0004597). Finally, JD and TB are kindly supported by the Czech Science Foundation JUNIOR STAR grant (No. 21-22435M), Czech Science Foundation grant (No. 22-30879S), Charles University PRIMUS grant (No. Primus/21/MED/003) and from Ministry of Education, Youth and Sports grant ERC CZ (LL2315).

