## Supplementary Figures for "Jag1 Insufficiency Disrupts Neonatal T Cell Differentiation and Impairs Hepatocyte Maturation, Leading to Altered Liver Fibrosis"

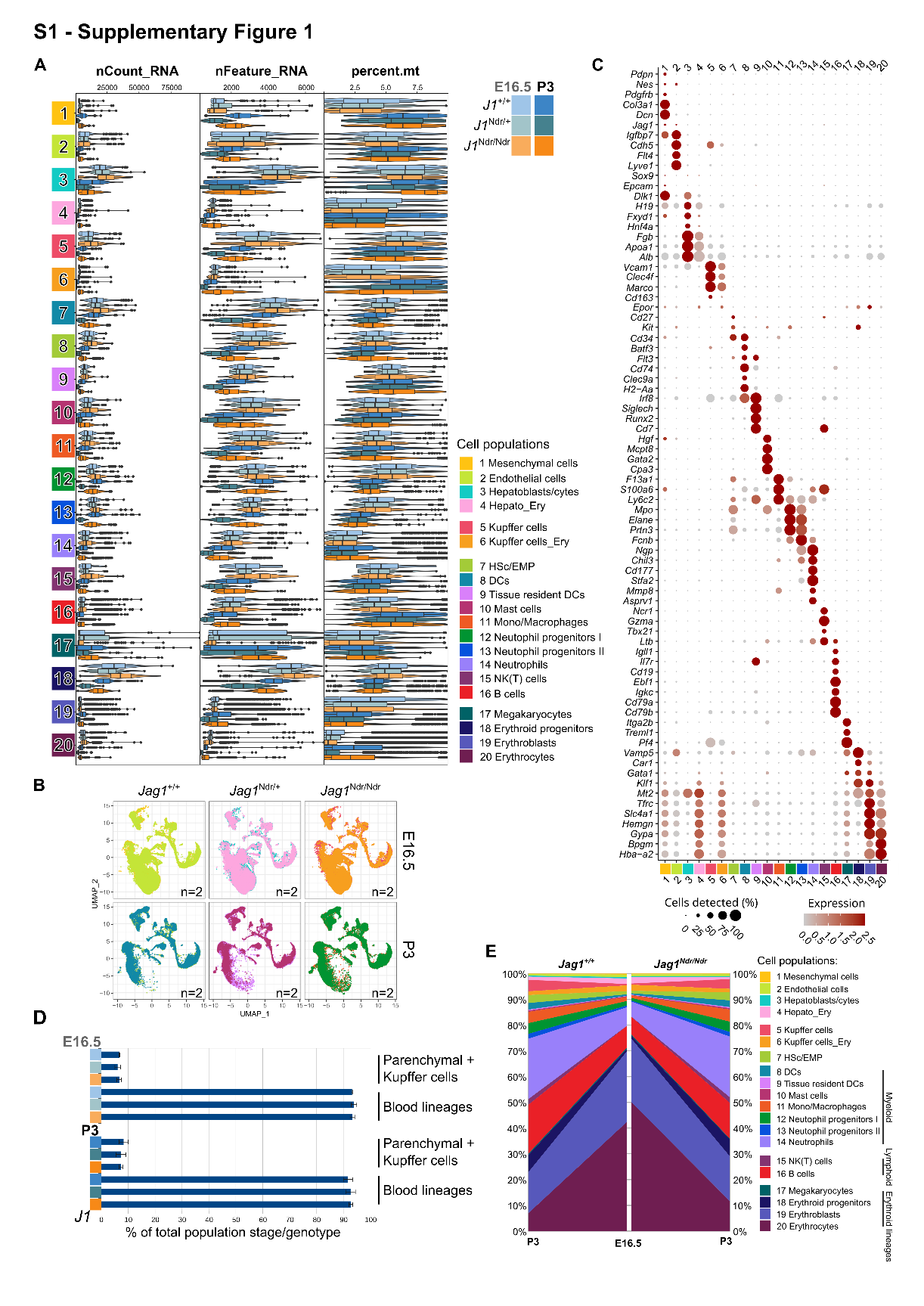
**Supplementary Figure 1**: ***Single-cell profiling of the developing and postnatal liver reveals a cell population shift from embryonic erythropoiesis to postnatal immune surveillance in Jag1^+/+^, Jag1^Ndr/+^, and Jag1^Ndr/Ndr^ mice.*** (**A**) Violin plot of the mitochondrial mRNA (percent-mt), unique gene counts (nFeature_RNA), and read counts (nCount_RNA) across the cell types identified in the *Jag1^+/+^*, *Jag1^Ndr/+^*, and *Jag1^Ndr/Ndr^* livers at E16.5 and P3. (**B**) Genotype, stage, or organ contribution to the composition of the scRNAseq liver samples including from E16.5 and P3 *Jag1^+/+^,* *Jag1^Ndr/+^,* and *Jag1^Ndr/Ndr^* mice. (**C**) Dot plot of the SCT-normalized mRNA marker expression of 78 DEGs characteristic for the individual cell types identified in B. (**D**) Bar graph representing the relative proportion of parenchymal and Kupffer cells (clusters 1-6) versus non-parenchymal cells (clusters 7-20) in dissociated *Jag1^+/+^*, *Jag1^Ndr/+^*, and *Jag1^Ndr/Ndr^* livers at E16.5 and P3. (**E**) Graph depicting relative cell type contribution in dissociated *Jag1^+/+^*, *Jag1^Ndr/+^*, and *Jag1^Ndr/Ndr^* livers at E16.5 and P3 normalized to the total sample cell count.

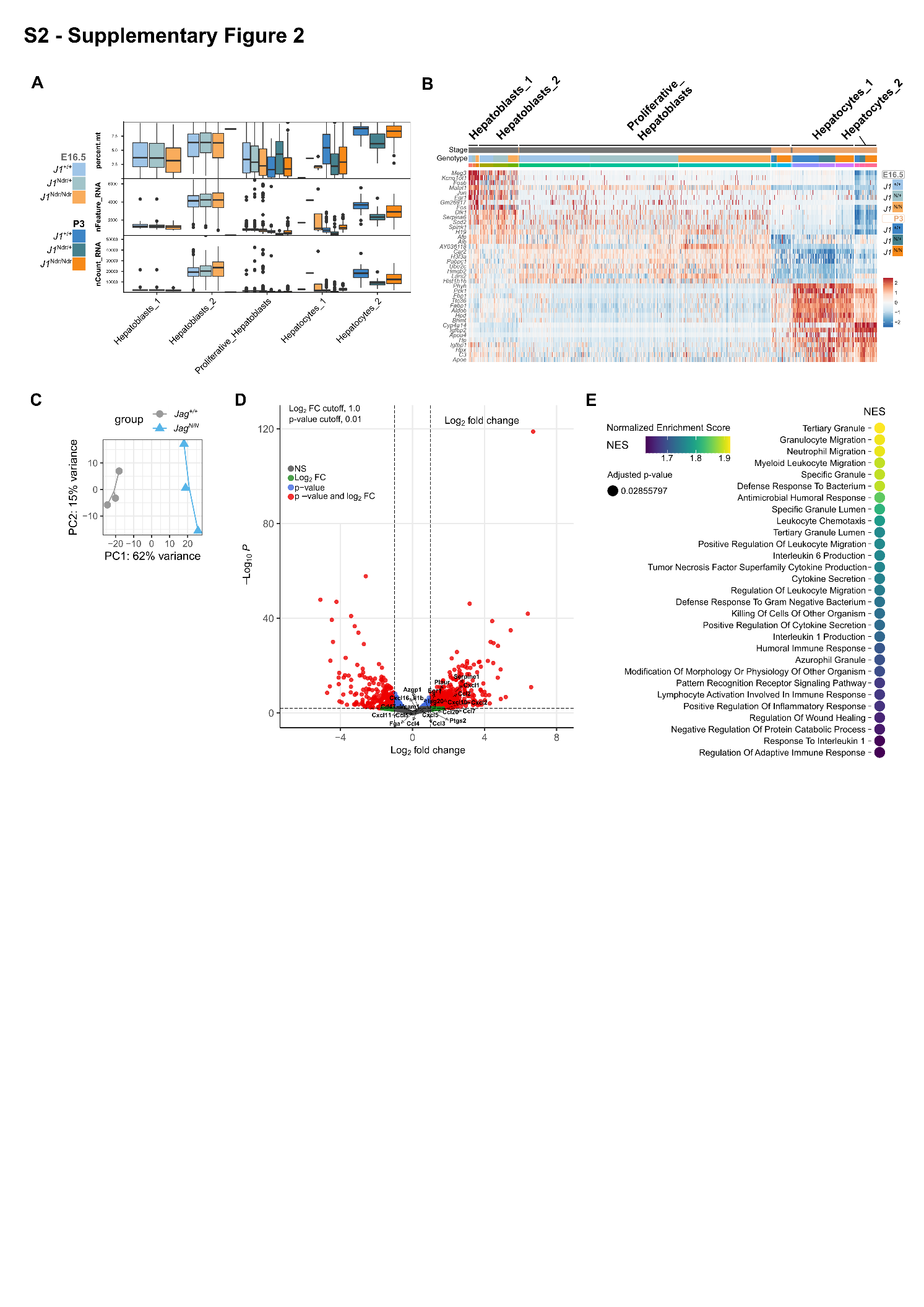
**Supplementary Figure 2**: ***Altered gene expression profile of Hepatocytes and whole livers in Jag1^Ndr/Ndr^ animals***. (**A**) Box and whiskers plot of the percentage mitochondrial mRNA (percent.mt), unique gene counts (nFeature_RNA), and read counts (nCount_RNA) across the hepatocyte-like cell types identified in the *Jag1^+/+^*, *Jag1^Ndr/+^*, and *Jag1^Ndr/Ndr^* livers by scRNA seq at E16.5 and P3. (**B)** Heatmap of the top 8 mRNA markers for each cluster. (**C**) Volcano plots of differentially expressed genes (DEGs) of *Jag1^Ndr/Ndr^* vs. *Jag1^Ndr/+^* and *Jag1^Ndr/Ndr^* vs. *Jag1^+/+^* Hepato_Ery and Hepatoblast/cytes at E16.5, and P3. (**C**) PCA distribution of the bulk RNAseq samples from Andersson et al., 2018, re-analyzed in Fig. 3 and Fig. S3F,G. (**D**) Volcano plot of the DEGs of *Jag1^Ndr/Ndr^* vs. *Jag1^+/+^* liver at P10. Pro-inflammatory markers of activated hepatocytes are highlighted. (**E**) Top29 GO pathways enriched in the *Jag1^Ndr/Ndr^* livers, NES (normalized enrichment score) indicates overlap of genes across the pathways.

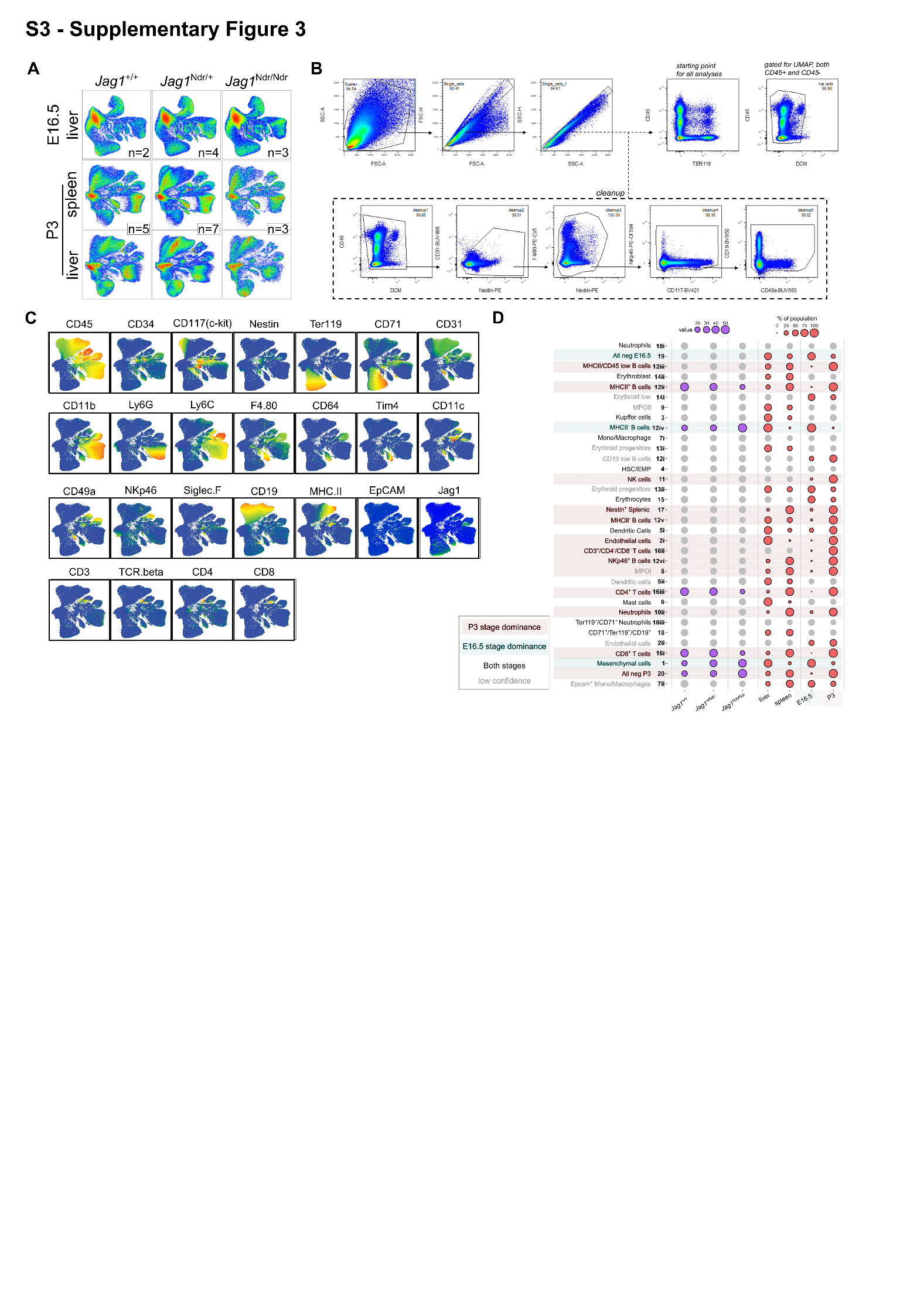
**Supplementary Figure 3**: ***25-colour Flow cytometry profiling of embryonic and postnatal liver and spleen reveals enrichment of B cells and depletion of T- cell populations in Jag1^Ndr/Ndr^ mice.*** (**A**) Genotype, stage, or organ contribution to the composition of the Flow cytometry spleen and/or liver samples from E16.5 and P3 *Jag1^+/+^,* *Jag1^Ndr/+^,* and *Jag1^Ndr/Ndr^* mice. (**B**) Flow cytometry gating strategy. (**C**) UMAP projections of the cells expressing marker proteins across the aggregated 25-colour Flow cytometry dataset of 194,999 cells sampled from E16.5 and P3 livers, and P3 spleens of *Jag1^+/+^*(n=7 biological replicates), *Jag1^Ndr/+^*(n=11 biological replicates), and *Jag1^Ndr/Ndr^* (n=6 biological replicates) mice (**D**) Dot plot depicting relative frequencies of the identified cell populations across genotypes, tissues, and developmental stages. MPO, myeloid progenitors; HSC, haematopoietic stem cells; EMP, erythroid-myeloid progenitor.

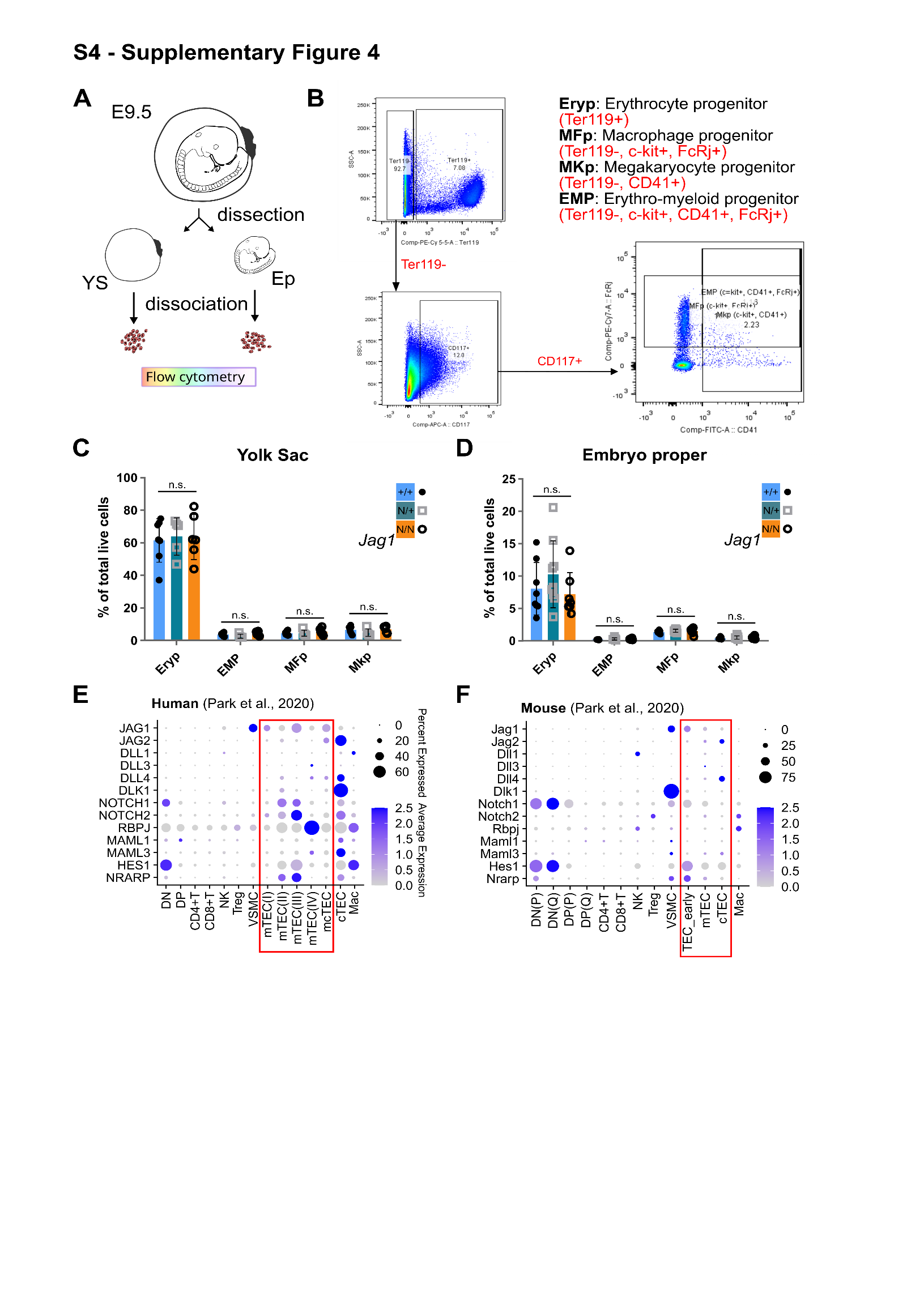

**Supplementary Figure 4**: ***The proportions of early embryonic haematopoietic progenitor populations are not altered in Jag1^Ndr/Ndr^ embryo proper (EP) and yolk sac (YS).*** (**A**) Schematic of the experiment. E9.5 embryos were harvested and yolk sac (YS) and embryo proper (EP) were processed separately for flow cytometry analysis. (**B**) Gating strategy for identification of the Erythrocyte progenitors (Eryp), Macrophage progenitors (MFp), Megakaryocyte progenitors (MKp), and Erythro-myeloid progenitors (EMP) after dead cell exclusion. (**C**, **D**) relative proportion of the live Eryp, EMP, MFp and MKp cells from *Jag1^Ndr/Ndr^* (n=7), *Jag1^Ndr/+^*(n=9), and *Jag1^+/+^* (n=8) yolk sac (C) and whole E9.5 embryos (D). (**E, F**) Dot plot of the reanalyzed median scaled ln-normalized mRNA expression of Notch signaling components in human (E) and mouse (F’) thymic cell populations from Park et al., 2020.; One-way ANOVA, multiple comparison with Bonferroni method.; n.s., no significant difference.

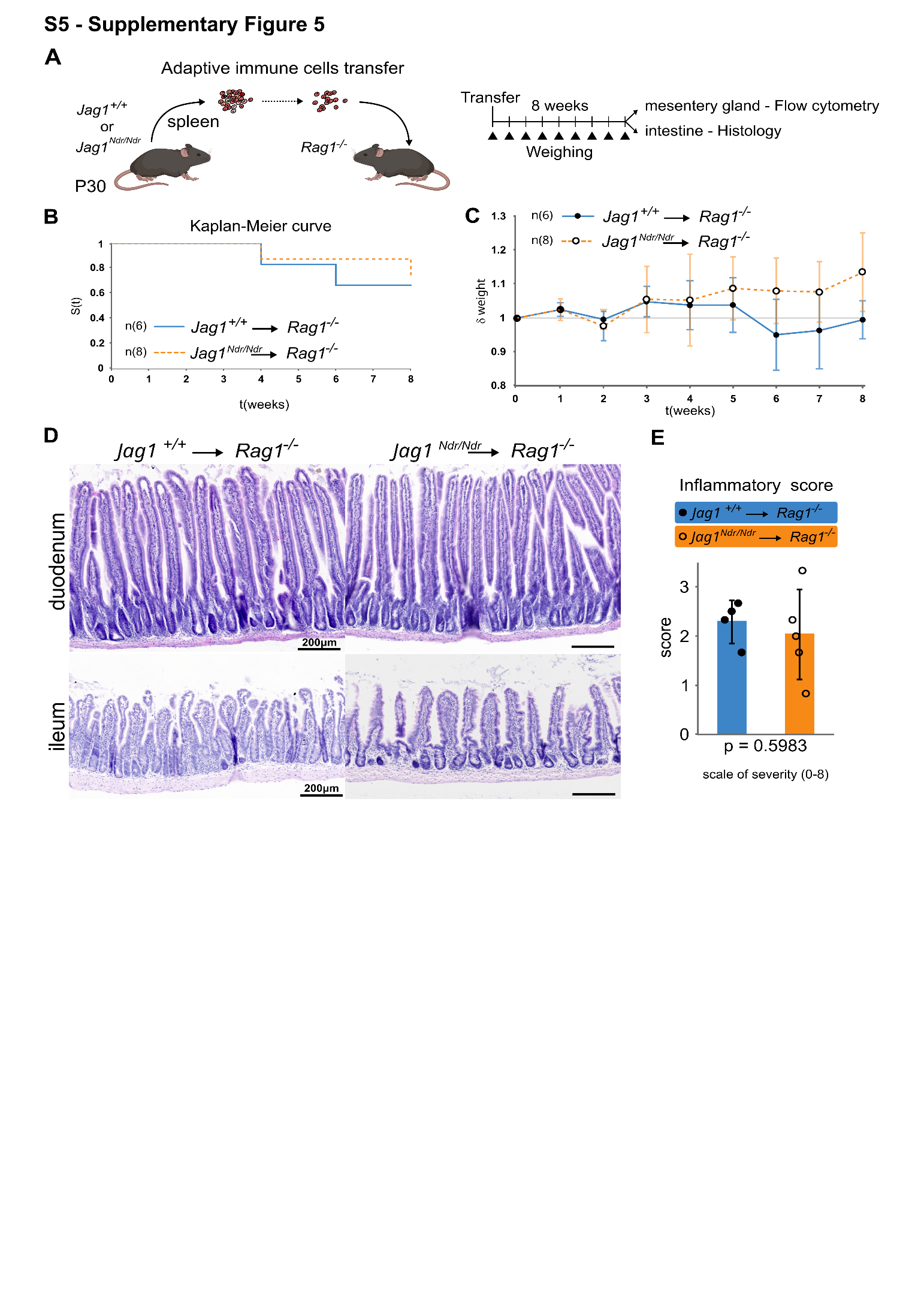
**Supplementary Figure 5**: ***Jag1^Ndr/Ndr^ T cells are not autoimmune as shown by the transfer to Rag1^-/-^ hosts. (A) Schematic of the experiment.*** (**B**) Survival curve of the *Rag1^-/-^* animals over the course of 8 weeks following transfer with *Jag1^+/+^* (n=6) or *Jag1^Ndr/Ndr^* (n=8) T cells. (**C**) Relative quantification of *Jag1^+/+^*→*Rag1^-/-^* and *Jag1^Ndr/Ndr^*→*Rag1^-/-^* mouse weight normalized to its original value on day 0 after T cell transfer over 8 weeks (1=100% of original weight, mean ± SD, n = 6-10 mice). (**D**) Representative H&E staining of intestinal sections (duodenum - top, ileum - bottom) performed 8 weeks after T cell transfer. (**E**) Comparison of the intestinal inflammatory score calculated using the H&E-stained slides of intestinal sections from the *Jag1^+/+^*→*Rag1^-/-^* (n=4) and *Jag1^Ndr/Ndr^*→*Rag1^-/-^* (n=5) mice. (mean ± SD, Statistical analysis was performed by unpaired, two-tailed Student’s t-test).

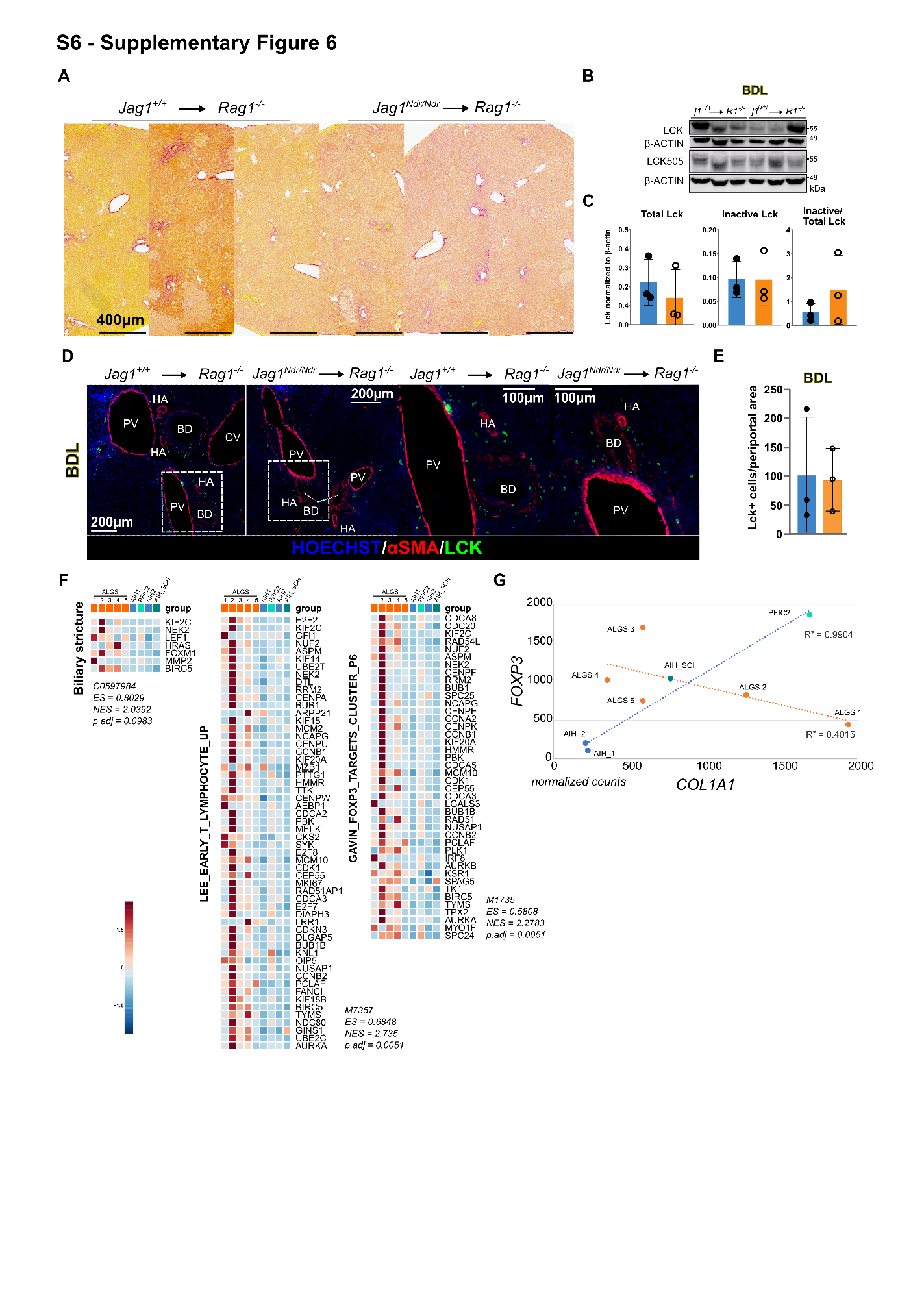

**Supplementary Figure 6**: ***Jag1^Ndr/Ndr^ T cells are present in periportal area after BDL treatment*** *(****A****)* *Representative Sirius red staining in Zone 3 of Jag1^+/+^→Rag1^-/-^ and Jag1^Ndr/Ndr^→Rag1^-/-^ mice after BDL.* (**B,C**) Western blot of total LCK, inactive Tyr505 LCK and β-ACTIN levels in LLL lysates from *Jag1^+/+^→Rag1^-/-^* and *Jag1^Ndr/Ndr^*→*Rag1^-/-^* mice after BDL (B) and respective quantification (C). (**D,E**) Representative immunofluorescent images of cryosections from the left lateral lobe of *Jag1^+/+^*→*Rag1^-/-^* (left) and *Jag1^Ndr/Ndr^*→*Rag1^-/-^* (right) mice after BDL treatment, stained with antibodies against Lck and SMA (D), and respective quantification of Lck^+^ in the periportal area (E). (**F**) Heat map of gene sets over- and under-represented in ALGS and control patients. (**G**) Correlation of *FOXP3* and *COL1A1* mRNA expression in patients with ALGS and control patients. CV, central vein; PV, portal vein; BD, bile duct; HA, hepatic artery AIH, autoimmune hepatitis; SCH, sclerosing cholangitis; PFIC2, Progressive familial intrahepatic cholestasis type 2; (N)ES, (normalized) enrichment score. Fisher Exact test was used for the gene set enrichment analysis; LLL, left lateral lobe; CV, central vein; PV, portal vein; BD, bile duct; HA, hepatic artery.
