## Supplementary Methods for "Jag1 Insufficiency Disrupts Neonatal T Cell Differentiation and Impairs Hepatocyte Maturation, Leading to Altered Liver Fibrosis"

***Mouse maintenance and breeding***

All animal experiments were performed in accordance with ARRIVE guidelines, local rules and regulations and all experiments were approved by Stockholm’s Norra Djurförsöksetiska nämnd (Stockholm animal research ethics board, ethics approval numbers: N50/14, N61/16, N5253/19, N2987/20) or Czech Academy of Sciences ethics board (approval number: AVCR 8362-2021). The *Jag1^Ndr/Ndr^* mice were maintained in a mixed C3H/C57bl6 genetic background as reported previously (Andersson *et al*, 2018) and are deposited in the European Mouse Mutant Archive EMMA: <https://www.infrafrontier.eu/search?keyword=EM:13207>. *Rag1*-deficient animals were purchased from Jax (Jax: 002216). Animals were maintained with standard day/night cycles, provided with food and water ad libitum, and were housed in cages with enrichment. Males and females were bred overnight and noon of day of plug was considered embryonic day (E) 0.5. Nodder mice were genotyped by the Transnetyx® (USA) automated qPCR genotyping company or by TaqMan qPCR assay.

***Adaptive T-cell transfer to Rag-deficient animals and DSS-induced colitis mouse model.***

Adult Rag1-deficient animals (Jax: 002216) were used as the acceptors of T and B lymphocytes isolated from *Jag1^+/+^* or *Jag1^Ndr/Ndr^* mice. Spleens from *Jag1^+/+^* or *Jag1^Ndr/Ndr^* mice (littermates) were passed through 40 um nylon mesh to obtain a single cell suspension, which was then incubated overnight in 10% complete RPMI (Sigma-Aldrich, #R8758) to remove adherent cells. After the incubation, the floating fraction containing mainly lymphocytes was collected. One million lymphocytes were injected into the recipient via tail vein, in PBS. For the induction of colitis by DSS (Sigma-Aldrich, # D8906) based on the(Chassaing *et al*, 2014), mice were given 2.5% DSS in drinking water for one week, followed by 10 days of recovery and subsequently one more week of DSS treatment. Scoring of the intestinal inflammatory status was performed on formalin-fixed, paraffin-embedded liver sections (4 μm) stained with hematoxylin and eosin (H&E) was assessed as described in (Erben *et al*, 2014).

***Bile duct ligation model of cholestasis***

The common bile duct ligation (BDL)-induced cholestais in mice was performed as in (Tag *et al*, 2015) with the following modifications. Mice were anesthetized with ketamine (80 mg/kg) and xylazine (10 mg/kg) and the extrahepatic common bile duct was cut in between the sutures. Mice were sacrificed after 7 days for further analyses.

**Immunohistochemistry and hematoxylin, and Sirius red staining**

Formalin-fixed, paraffin-embedded liver sections (4 μm) were stained with hematoxylin and eosin (H&E) or Sirius Red (SR)(Chalupský *et al*, 2013). For immunofluorescence, paraffin sections were subjected to heat-induced antigen retrieval in citrate (pH 6) buffer, and incubated with primary antibodies (**Methods Table 1**) overnight at 4°C or 1h/37°C. Secondary antibodies were incubated for 120 min at 22°C

**Liver section image acquisition**

Mice > P10 were euthanized with CO2 or < P10 were decapitated, and liver was immediately dissected out, left lateral lobe was cut in half, one embedded in OCT and frozen on dry ice, and other fixed O/N in 4% PFA (VWR, #20909.290) in PBS,and processed for paraffine sectioning. Immunofluorescent staining was carried out on cryosections (14 µm) or paraffine sections (5 µm) as previously described (Hankeova *et al*, 2021; Andersson *et al*, 2018). The liver was imaged using the LSM 980 confocal microscope or Axioscan (Carl Zeiss).

Fresh frozen cryosections from left lateral lobe were postfixed in 4% formaldehyde for 10 minutes, washed 3x in PBS, blocked and permeabilized in PBS containing 10% BSA and 0.3% TritonX-100 for 1 hour at room temperature (RT). Tissue sections were then incubated with primary antibodies in blocking solution overnight at 4°C. Next, the sections were washed 3x with PBS, and incubated with fluorochrome-conjugated secondary antibody for 1 hour at RT. Following the incubation, the sections were washed 3x with PBS, and incubated with Hoescht 3342 for 10 minutes at RT and washed again 3x with PBS and mounted in MOWIOL and imaged with the Axioscan (Carl Zeiss).

SR-positive collagen deposits were measured from scanned whole liver sections (scanned using slide scanner Axio Scan.Z1, Carl Zeiss MicroImaging, Jena, Germany) and quantified by Fiji software. Keratin 19-, aSMA- and collagen-positive (collagen+) areas were measured from portal field (PF) view obtained with 40x objective (HCX PL APO 40x/0.75) from liver sections and quantified in Fiji software using IJ_IsoData (collagen), RenyiEntropy (aSMA), or IJ-IsoData (K19) thresholds.

For Alfa-fetoprotein and Cyp1A2 on paraffin sections as described in (Andersson *et al*, 2018). For AFP immunostaining, the slides were treated with quenching agent TrueView (SP-8500-15, Vector laboratories) to remove non-specific background signal. Slides were mounted with Dako Fluorescent mounting medium (S3023) and imaged on Nikon / CrEST X-Light V3 Spinning Disk confocal microscope.

**Western blotting**

Mouse liver tissue was weighed and homogenized in RIPA Lysis and Extraction Buffer (Thermo Scientific, #89900) with the Protease and phosphatase inhibitor (Thermo Scientific, #A32961), using 15-25mg of tissue/200μl. Tissue samples were disrupted using a chilled glass-pestle homogenizer. Samples were then agitated for 2 hours at 4°C, and subsequently centrifuged at 16,000 rcf for 20 min at 4°C. The supernatant was collected, and the protein concentrations determined using the Pierce™ BCA Protein Assay Kit (Thermo Scientific, #23225). For western blotting, aliquots of 25 μg were denatured by boiling in Laemmli Sample Buffer (2x), separated by SDS-PAGE, and transferred onto PVDF membranes (Amersham™ Hybond®, # *GE10600021*) by electroblotting. The membrane was cut, blocked by 5% non-fat milk in 1X TBST (Tris-buffered saline, 0.1% Tween 20) for 1h/RT, and incubated ON/4°C with a Rabbit anti-Lck, and Mouse anti-β-actin antibodies. The following day, membranes were washed 4x/10min in 1X TBST and incubated with corresponding HRP-conjugated secondary antibodies for 1h/RT, followed by 4x/10min wash in 1X TBST. Chemiluminescence was detection with Pierce™ ECL Western Blotting Substrate (Thermo Fisher Scientific, #32209) on ChemiDoc machine (Bio-Rad).

**Flow cytometry: E16.5 and P3 sample preparation and sequencing**

Reagents for the single-cell isolation: Tabletop centrifuge with cooling; QuadroMACS separator (Miltenyi, *#*130-091-051); Dead Cell Removal Kit (Miltenyi, #130-090-101); *sterile 5ml Transfer Pipettes (Merc, #HS206371C-500EA);* DPBS -Mg^2+^,-Ca^2+^ (Gibco, #*D8537*); HBSS -Mg^2+^,-Ca^2+^ (Gibco, #*14175095*); 2ml Eppendorf DNA LoBind® tubes (Merck, #*EP0030108078*), Liver digest medium (LDM) (Gibco; #17703034) - unfreeze in 4C, protect from light, avoid freeze-thaw; TrypLE Express (Gibco, 12604013); Millex-GV Filter 0.22 µm (Merck-Millipore, #*SLGVR33RS)*; 40μm strainers (BD, #352350) or EASYstrainer (Greiner, #542040); 5ml flow cytometry tubes with 70μm filter cap (Falcon, #38030); FCS (Gibco, #*10270106*); 0.5M EDTA pH 8.0 (Thermo Scientific, #*R1021*); BSA (Sigma-Aldrich, #A7906).

Buffers:

E16.5 - Wash Buffer (WB16-E) 1% BSA in HBSS, filter sterile (0.22 µm), Wash Buffer (WB16) 1% BSA, 1mM EDTA, HBSS, filter sterile (0.22 µm).

P3 - Wash Buffer (WBP3-E) 1% FCS in HBSS, filter sterile (0.22 µm), Wash Buffer (WBP3) 1% FCS, 1mM EDTA, HBSS, filter sterile (0.22 µm).

Embryo stage was determined based on the presence of vaginal plug, the morning after breeding initiation, noon of day of plug was considered as E0.5. All embryos at E16.5 and P3 pups from the *Jag1^Ndr/+^* x *Jag1^Ndr/+^* breeding were sacrificed by decapitation, their livers (and spleens at P3) dissected out and placed in 12- or 24-well plates with ice-cold DPBS on ice unless stated otherwise. The sex of P3 pups was determined based on scrotum pigmentation (Wolterink-Donselaar *et al*, 2009).Material for genotyping was collected (tail or limb tissue) and processed for genotyping using TaqMan probe (Thermo Fisher) yielding results in 3 hours (P3 Jag1^Ndr/Ndr^ pups were identified based on the presence of jaundice and random Jag1^Ndr/+^ and Jag1^+/+^ controls genotypes were confirmed afterwards). The tissue was kept in ice-cold DPBS for 30min before being transferred into 5ml flow cytometry tubes with 1.5 ml LDM + 0.5 ml of TrypLE Express (E16.5) or 2ml LDM (P3 stage), pipetted 5x through sterile wide-bore blue tip (E16.5) or transfer pipettes (P3) and incubated at 37°C/5min in a water bath (inverting each tube after 2-3 minutes). After the 5 min incubation, the tissue was disrupted further by gentle pipetting 10x using a filtered 1 ml tip, followed by an additional round of incubation at 37°C/5min in a water bath (inverting each tube after 2-3 minutes). The enzymatic digestion was terminated by addition of 1.5 ml of WB16-E/ WBP3-E and additional mechanical disruption of the remaining tissue by pipetting each sample 20x using filtered 1 ml wide-bore blue pipette tip. We immediately poured the suspension through 70 µm filters into flow cytometry tubes, rinsed the filter once with 0.5 ml WB16/WBP3, and passed the filtered suspension through a 40 µm strainer, each sample into two 2 ml Eppi tubes.

The duplicates of E16.5/P3 samples were centrifuged at 100g for 5mins at 5°C. Each supernatant was transferred to two new tubes, while the pellets were resuspended in 200 μl of Dead Cell Removal MicroBead suspension in Binding buffer and the resuspended pellets were further incubated for 5min at RT. In the meantime, the supernatant from the previous step was centrifuged at 250g for 5min at 5°C. The new supernatant was discarded, and the corresponding pellets were mixed with 200 μl of Dead Cell Removal MicroBeads with the previous cell suspension in Binding buffer. The mixture was incubated for 10min at RT. After the incubation 300 μl 1× Binding Buffer was added to the cell suspension to obtain a final volume of 500 μl, suitable for magnetic separation with MS columns. The separation was done based on the manufacturer’s recommendations. Briefly, we rinsed the columns with 500 μl of 1× Binding Buffer, applied the cell suspension onto the column, and collected the flowthrough containing unlabeled cells (living cells). Columns were further washed three more times with 500μl of 1× Binding Buffer, flowthrough collected and 20 μl filter sterile 10% BSA was added to the purified cells. Resulting samples were divided in two, whereof 250 μl was used for scRNAseq, following the 10x Genomics® Single Cell Protocol, and the remainder was used for the flow cytometry as described below.

The duplicate P3 samples were centrifuged at 100g for 5min at 5°C, and supernatants were transferred to new tubes. The pelleted cells from each duplicate were pooled and resuspended in 1.5 ml of the WBP3-E buffer and transferred to a new 2 ml tube. In the meantime, the supernatants from the previous step were centrifuged at 250g for 5min at 5°C. The supernatants were discarded, and the corresponding pelleted cells were resuspended in 0.5 ml of the WBP3-E buffer and added to the resuspended cells obtained from the first centrifugation. The full sample cell suspensions (now in 2 ml of WBP3-E buffer) were then transferred into a new protein lo-bind tube through 40 μl cell strainer to remove any remaining cell debris or large clumps. We then determined the cell concentration using a Countess® II Quantification to calculate the appropriate volume for the subsequent resuspension to obtain the target concentrations of ~1200cells/μl as recommended by the 10x Genomics® Single Cell Protocol. The whole procedure including tissue collection, genotyping and dissociation took 3 hours, before initiation of processing with 10x microfluidics chromium and/or for flow cytometry.

**Library preparation and sequencing**

The E16.5, P3 samples were processed on a Chromium microfluidics platform (10X Genomics) using the Chromium^TM^ Single Cell 3’ Library and Gel Bead Kit v2 (10X Genomics, # PN-120237) and the Chromium^TM^ Single Cell A Chip Kit (10X Genomics, #PN-120236) following the manufacturer’s instructions. The E16.5 library preparation, validation with Bioanalyzer High Sensitivity DNA kit (Agilent, # 5067-4626), and sequencing with NovaSeq (100 cycles) platform was performed by the SciLifeLab, Stockholm. The P3 samples, library preparation, validation with Bioanalyzer High Sensitivity DNA kit (Agilent, # 5067-4626), and sequencing with NextSeq550 (75 cycles) or S4 Novaseq platform was performed by the *Bioinformatics and Expression analysis core facility (*BEA), Huddinge, Stockholm. All scRANseq data are available online (GSE236483).

**Analysis of the scRNA seq using Seurat packages**

Demultiplexing, quality control, raw data, gene expression data counts were performed using CellRanger (Chromium), by the BEA and SciLife core facilities. Matrices were further processed and analyzed with R (version 4.1.2) in R Studio. Prior to transformation into Seurat Objects, matrices were precleared using SoupX (Young & Behjati, 2020) with default settings of setContaminationFraction parameter (sc) for the P3, and (sc, 0.13) for the E16.5 stages. All bioinformatic analyses was performed using Seurat pipeline, doublet removal with scDblFinder (Germain *et al*, 2022), QC cutoffs for E16.5 datasets were nFeature_RNA > 500 & nFeature_RNA < 7000 & percent.mt < 10, for P3 a range of nFeature_RNA > 300-500 & nFeature_RNA < 4500-6000 & percent.mt < 10 were used, followed by SCT based integration of the E16.5 and P3 Jag1^+/+^, Jag1^Ndr/+^, and Jag1^Ndr/ Ndr^ datasets (Butler *et al*, 2018). The high prevalence of erythrocytes and erythroblasts in embryonic/neonatal liver may have contributed ambient RNA to the droplet-based scRNA-seq: despite *in silico* free-mRNA decontamination and doublets removal, erythrocyte signatures persisted in a subset of hepatoblasts/hepatocytes and Kupffer cells. In total, 183,542 cells passed through the cross-contamination adjustment, quality control filtering and doublet removal in each of the duplicates per genotype and stage, analyzed in Seurat (Butler *et al*, 2018) (**Fig. 1A,B, S1A-C**). A mean of 1520 genes, 6703 reads, and 3.0% of mitochondrial mRNA content were detected per cell*.* For comparative transcriptomic analysis, logNorm and scaled RNA data were used. Liver parenchymal cells constituted ~6.5% of cells at E16.5, and ~7.5% of cells at P3 and included mesenchymal cells, endothelial cells, hepatoblasts and hepatocytes (**Fig. S1D**), this parenchymal proportion is lower than *in vivo*, but consistent with *ex vivo* liver digest (Guilliams *et al*, 2022). The persistent cross contamination was in the subset Hepatoblast/Hepatocyte and Hepatocyte_Ery cells removed by subtracting the top 40 marker *mRNA* of Erythroblasts, Erythrocytes and B cells (Supplementary Table, **ST9**). For the hepatoblasts/hepatocyte DGE analysis (**ST1-5**) we analyzed each cluster separately using the Seurat FindMarkers function.

**Deconvolution of bulk RNA seq using MuSiC packages**

To estimate cell type proportions in *Jag1^Ndr/Ndr^* livers compared to *Jag1^+/+^* livers at P10, we performed cell type deconvolution of our previously generated bulk RNA-seq datasets GSE104875 (Andersson *et al*, 2018) employing MuSiC package for Multi-subject Single-cell Deconvolution (Wang *et al*, 2019) that uses single cell RNA-seq data as a reference for deconvolution.

Deconvolution using an annotated scRNAseq dataset available under GEO accession number GSE171993 (Liang *et al*, 2022): The dataset was imported into R 4.1.0 package Seurat 4.0.5 (Hao *et al*, 2021) and stages P1, P3, and P7 were subset. The original annotation accounted for 30 cell types in liver scRNA-seq data (Liang *et al*, 2022). We filtered for high-quality cells containing < 5% of mitochondrial genes expressed per cell and 200 - 4000 genes per cell, and for genes that are expressed in at least 3 cells, resulting in a dataset of 32,967 cells and 23,170 genes. The Seurat object was converted to an Expression set, required as input in MuSiC using a function SeuratToExpressionSet from BisqueRNA 1.0.5 R package (Jew *et al*, 2020). The MuSiC was performed using music_prop function. To analyze the differentiation signature of hepatocytes, we further subset neonatal hepatocytes (P1, P3, P7) from scRNA-seq dataset (1281 cells), converted to Expression set and used this as a scRNA-seq reference for MuSiC on the P10 bulk RNA-seq dataset.

Deconvolution using a time-course mSTRT scRNA-seq dataset on isolated albumin-positive cells (GSE209749) (Yang *et al*, 2023): The analysis was done in R version 4.3.0 with updated MuSiC v1.0.0 that supports SingleCellExperiment class as single cell reference. The scRNA-seq count matrix “GSE209749_readcount.mSTRT-seq.csv.gz” was downloaded from GEO and loaded into R. The time-course dataset was subset (removing P60_2N, P60_4N and P60_8N cells belonging to a different experiment) and basic quality control was performed with SingleCellExperiment (Amezquita *et al*, 2020), scater (McCarthy *et al*, 2017) and scran packages(Lun *et al*, 2016). Cells with low library size and low number of detected genes were filtered out, leaving 1783 cells in the time-course dataset stored as SingleCellExperiment object. Estimated proportions of different stages (E17.5 – P60) were generated with *music_prop* function applied on P10 bulk RNAseq matrix.

***Gene set enrichment analysis (GSEA) for a hepatoblast or hepatocyte signatures***

To derive hepatoblast and hepatocyte gene expression signatures we used GEO bulk RNA-dataset dataset GSE176069. In the dataset, primary Dlk1^+^ enriched hepatoblasts from E14.5 livers and primary hepatocytes from adult livers of C57BL/6JOlaHsd (Harlan Laboratories/Envigo) or C57BL6/JRj (Janvier Labs) mice (Belicova *et al*, 2021). The provided raw counts matrix was analyzed in R (version 4.1.0) with DESeq2 (Love *et al*, 2014)(version 1.34.0) package to identify differentially expressed genes using Wald test. The genes were sorted by p-adjusted value (default Benjamini and Hochberg method) and log2 fold change. The top 500 most significant genes were considered markers of hepatoblasts and hepatocytes, respectively.

The top 500 markers were provided as gene lists to GSEA desktop tool (version 4.3.2, Broad Institute) to analyse whether there is an enrichment of heptoblasts or hepatocyte signature in our previously generated bulk RNA-seq datasets GSE104875, and GSE104873)(Andersson *et al*, 2018) on P10 *Jag1^+/+^* and *Jag1^Ndr/Ndr^* livers, and control and ALGS patients, respectively. These datasets were processed with DESeq2 from raw counts to size factor normalized counts as input for the GSEA desktop tool. The genes were ranked based on Signal2Noise and permutation was done using the “gene_set” parameter. The results were replotted using ggplot2 (version 3.4.0) package in R. The top 50 enriched genes per signature were visualized using heatmaps.

***Bulk RNA Sequencing***

Files from GSE104875 and GSE104875 series were downloaded and converted to fastq files using SRA Toolkit (NCBI) and aligned using STAR aligner tool(Dobin *et al*, 2013) to the genomes of *Mus musculus* (GRCm38.p6) and *Homo sapiens* (GRCh38.p13) respectively (downloaded from NCBI). Reads aligning to gene exons were counted using featureCounts program from Subread package(Liao *et al*, 2014) Out of these, lists of differentially expressed genes (DEGs) between groups as well as normalized read counts were obtained using DESeq2 package(Love *et al*, 2014). Considering the study design, all genes showing p-value (adjusted by Benjamini-Hochberg method) < 0.05 were further examined for the enrichment of functional processes. To identify differentially expressed pathways, gene list of DEGs were queried by clusterProfiler package(Yu *et al*, 2012) against Disease Ontology and MSigDB databases using overrepresentation DisGeNet and GSEA analysis.

**Sample processing and flow cytometric analysis of embryo proper and yolk sac at E9.5**

For dissociation and flow cytometry we followed a protocol described by Balounová and colleagues (Balounová *et al*, 2019). In brief, E9.5 embryo proper (EP) and yolk sac (YS) were dissected and dissociated separately. After briefly washing in cold Hank’s balanced saline solution (HBSS) (Gibco, #14175095). EP and YS were incubated with 1 mg/mL Dispase (Gibco, #17105041) in HBSS for (~ 10 min for EP, ~17 min for YS) in 1.5mL Eppendorf tubes in water bath at 37°C, and occasionally mixed by gentle pipetting. The reaction was stopped by washing in HBSS with 2% FCS (Gibco, #10270106). Suspensions were passed through a 40μm cell strainer (BD, #352350) and centrifuged at 350g/7min/4°C. Pelleted cells were resuspended in 200ul flow cytometry wash (DPBS with 2% FBS, 2mM EDTA, Thermo Fisher, #AM9262) and transferred to 96-well V-bottom plates (Sigma-Aldrich, #BR781601) for further flow cytometric analysis. Primary antibody mixes (**Methods Table 2**) were added and incubated for 30 min at 4°C, followed by LIVE/DEAD Fixable Aqua Dead Cell Stain (Thermo Fisher, # L34957) incubation for additional 30 min at 4°C. Cells were fixed for 15min/RT with fixation buffer from eBioscience™ FOXP3/ Transcription Factor Staining Buffer Set (eBioscience, #00-5523-00). Acquisition was performed on LSRII flow cytometer (BD Biosciences). All data were analysed using FlowJo software (FlowJo, V10.7.1).

**Flow cytometry E16.5, P3 - *Staining***

The flow cytometry was performed based on (Filipovic et al, 2019)(Filipovic et al, 2019). Single cell suspensions (cells) from liver and spleen were washed twice and resuspended in flow cytometry buffer (PBS with 2 mM EDTA and 2% FBS), filtered through a 100 µm strainer (BD Falcon) and stained in 96-well V-bottom plates. Unless otherwise stated, staining steps were performed with antibodies diluted according to the table X in 50 µl of the flow cytometry buffer, at 4ºC in the dark, and washing steps were performed by resuspending the sample in 150 µl of the flow cytometry buffer and centrifuging plates for 5 minutes at 500g at 4ºC. The BD Horizon Brilliant Stain Buffer Plus was added to the antibody mix in every step when the BD Horizon Brilliant dyes were used. Cells were pre-incubated with TruStain FcX (anti-CD16/32) to block Fc receptors for 10 minutes at room temperature. Cells were first stained with anti-CD64 antibody for 45 minutes. Next, cells were stained with antibodies against other surface antigens (**Methods Table 3**) for 30 minutes, followed by two washes in flow cytometry buffer. Cells were then stained with the LIVE/DEAD Fixable Aqua Dead Cell Stain (Thermo Fisher) and fluorescently conjugated streptavidin for 30 minutes. This was followed by two washes in flow cytometry buffer. Next, cells were fixed for 45 minutes in 100 µl fixation/permeabilization working solution from eBioscience Foxp3/Transcription Factor Staining Buffer kit (Thermo Fisher) at room temperature. After this step, cells were washed in 1x permeabilization buffer from the same kit. Finally, cells were stained with antibodies against intracellular antigens diluted in 1x permeabilization buffer from the same kit for 30 minutes at room temperature. Samples were then washed twice in 1x permeabilization buffer and resuspended in flow cytometry buffer for acquisition. For single-stained compensation controls, UltraComp eBeads Compensation Beads (Thermo Fisher) were used according to manufacturer’s instructions. FACSymphony A5 flow cytometer (BD Biosciences) was used for acquisition. The instrument was equipped with the following lasers: UV (355 nm), violet (405 nm), blue (488 nm), yellow/green (561 nm) and red (637 nm). See Table 3 for details of the panel and cytometer configuration settings.

**Flow cytometry E16.5, P3 - Analysis**

FCS3.0 files were exported from the FACSDiva and imported into FlowJo v.10.7.1 for analysis. The following plugin versions (from FlowJo Exchange) were used: FlowAI (2.1), DownSample (3.2), UMAP (3.1), PhenoGraph (3.0). The data were pre-processed using FlowAI (parameters selected: all checks, second fraction FR = 0.1, alpha FR = 0.01, maximum changepoints = 3, changepoint penalty = 500, dynamic range check side = both) to remove any anomalies present in FCS files. A compensation matrix was generated using AutoSpill in FlowJo. Events were downsampled from the live cell gate from all samples using DownSample, and categorical values were added to the downsampled populations (e.g., genotype, tissue) prior to concatenation so that groups of interest could be deconvoluted during the analysis. UMAP and PhenoGraph were run using all parameters from the panel except BV510. As over- and under-represented input groups are similarly weighted in the PhenoGraph output clusters, PhenoGraph results were normalized to account for the total number of cells from each input group. Some figures were generated in RStudio (version 1.3.959) using RColorBrewer (v1.1-2), ggplot2 (v3.2.1 and v3.3.0), tidyr (v.1.0.2), reshape2 (v.1.4.3), and pheatmap (v.10.12).

***Flow cytometry analysis of thymic epithelial cells***

Thymi from 4-week-old *Jag1^+/+^* or *Jag1^Ndr/Ndr^* mice were minced by scissors into small pieces and dissociated by enzymatic digestion for 30min at 37 °C using 0.3 mg ml−1 collagenase D (Roche), 1 mg ml−1 dispase II (Gibco) and 10 ng ml−1 DNase I (Sigma-Aldrich) in RPMI supplemented with 2% FBS. Percoll density gradient centrifugation was performed to enrich for thymic epithelial cells. Cells were resuspended in 2 ml of 1.115 g ml−1 isotonic Percoll (Sigma-Aldrich) and placed at the bottom of a tube. Subsequently, 1 ml of isotonic 1.065 g ml−1 Percoll and then 1 ml of PBS were layered on top. The Percoll gradient was run at 2,700 rpm (1451xg), at 4 °C, with no acceleration or brake for 30 min in the Eppendorf 5804 R centrifuge with S–4–72 rotor. The thymic epithelial cells were collected and subjected to flow cytometric analysis using the following set of antibodies for extracellular staining: CD45 APC-Cy7, EpCAM, APC, Ly6d PB, MHCII PE (all Biolegend). Intracellular staining of Aire AF488 (Invitrogen) was done after fixation and permeabilization of cells by the Foxp3/Transcription factor fixation buffer set (Thermo) according to manufacturer’s instructions. Extracellular cell staining for flow cytometry analysis was performed at 4°C, in the dark, on ice, for 30 minutes, while the fixation and intracellular staining was performed at room temperature, in the dark for 40 minutes (each step). Dead cells were excluded by addition of Viability dye 506 (Thermo) prior to fixation. Samples were analysed using a BD LSRII flow cytometer and FlowJO software version 10.7 (BD).

***Flow cytometry analysis of the T-cell compartment***

Spleens and thymi of *Jag1^+/+^* or *Jag1^Ndr/Ndr^* mice were passed through 40 um nylon mesh to obtain a single cell suspension. The following antibodies were used for T-cell analysis: CD4 BV785, CD44 BV 711, CD8 Percp-Cy5.5, TCR-B APC-Cy7 (all Biolegend). Extracellular cell staining for flow cytometry analysis was performed at 4°C, in the dark, on ice, for 30 min. Dead cells were excluded from analysis by addition of Viability dye 506 (Thermo) prior to the fixation. The fixation was performed at room-temperature in dark for 40 minutes using Foxp3/Transcription factor fixation buffer set (Thermo). Foxp3 BV421 (BD) staining was done at room temperature in dark for 40 minutes. Samples were analysed using BD LSRII flow cytometer and FlowJO software version 10.7 (BD).

**Supplementary Methods Table 1. Antibodies used in immunohistochemical or western blot analysis.**

| **Antigen** | **Clone** | **Dilution used** | **Company** |
| --- | --- | --- | --- |
| anti-mouse-HRP | #ab205719 | WB 1:20000 | Abcam |
| anti-rabbit-HRP | #ab205718 | WB 1:20000 | Abcam |
| Donkey anti-Rat IgG (H+L) Highly Cross-Adsorbed (also Ms) Secondary Antibody, Alexa Fluor™ 647 | A78947 | IF 1:250 (1:500 for the FOXP3,ECAD,LCK staining) | ThermoFisher |
| anti-rabbit-AlexaFluor-488 | 711-545-152 | IF 1:250 | Jackson ImmunoResearch |
| anti-mouse-RhodamineRedX | 715-295-150 | IF 1:250 | Jackson ImmunoResearch |
| anti-β-actin | #sc-47778 | WB 1:2000 | Santa Cruz Biotechnology |
| Lck | n.a.(Veillette et al, 1988)(Veillette et al, 1988) | WB/IF 1:1000 | Kind gift from Dominik Filipp |
| Tyr505Lck | #2751 | WB 1:1000 | Cell Signaling |
| aSMA – paraffine sections | M0851 | IF 1:50 | Dako |
| aSMA - cryosections | A2547 | IF 1:500 | Sigma-Aldrich |
| Collagen I- paraffine sections | ab21286 | IF 1:100 | Abcam |
| ECAD - cryosections | 610181 | IF 1:100 | BD biosciences |
| FOXP3 - cryosections | BV421 | IF 1:500 | BD |
| CK19- paraffine sections | TROMA-III RRID: AB_2133570 | IF 1:250 | DSHB |

**Supplementary Methods Table 2. Antibodies used in E9.5 flow cytometry.**

| **Antigen** | **Clone** | **Fluorophore** | **Laser line** | **BD LSRII** | **Dilution used** | **Company** |
| --- | --- | --- | --- | --- | --- | --- |
| CD41 | MWReg30 | BB515, FITC | Blue (488 nm) | 525/50 | 100 | Biolegend, #133903 |
| CD117 | ACK2 | APC/eF660 | Red (640nm) | 670/14 | 50 | Invitrogen, #17-1172-83 |
| Viability dye | N/A | BV510 | Violet  (405 nm) | 525/50 | 1000 | Thermo Fisher |
| CD45 | 30-F11 | Qdot585, BV570 |  | 585/42 | 100 | Biolegend, #103135 |
| TER119 | TER-119 | PE-Cy5.5 | Yellow-Green  (532nm) | 710/50 | 100 | Invitrogen, #35-5921-82 |
| FcRγ | 93 | PE-Cy7 |  | 780/60 | 50 | Biolegend, #101317 |

**Supplementary Methods Table 3. Antibodies used in 25-color flow cytometry.**

| **Antigen** | **Clone** | **Fluorophore** | **Laser line** | **BD FACSymphony filter** | **Dilution used** | **Company** |
| --- | --- | --- | --- | --- | --- | --- |
| CD45 | 30-F11 | BUV395 | UV  (355 nm) | 379/28 | 100 | BD Biosciences |
| CD31 | 390 | BUV496 |  | 515/30 | 100 | BD Biosciences |
| CD49a | Ha31/8 | BUV563 |  | 580/20 | 400 | BD Biosciences |
| TCR-beta | H57-597 | BUV615 |  | 605/20 | 100 | BD Biosciences |
| TER119 | TER-119 | BUV661 |  | 670/25 | 200 | BD Biosciences |
| CD11b | M1/70 | BUV737 |  | 735/30 | 200 | BD Biosciences |
| EpCAM | G8.8 | BUV805 |  | 810/40 | 100 | BD Biosciences |
| Ly6C | AL-21 | FITC | Blue  (488 nm) | 530/30 | 200 | BD Biosciences |
| CD34 | MEC14.7 | Biotin |  | N/A | 50 | Biolegend |
| Streptavidin | NA | BB630 |  | 610/20 | 400 | BD Biosciences |
| Tim-4 | RMT4-54 | BB700 |  | 710/50 | 100 | BD Biosciences |
| Jag1 | HMJ1-29 | APC | Red  (637 nm) | 670/30 | 100 | Biolegend |
| CD3 | 17A2 | AF700 |  | 730/45 | 200 | Biolegend |
| MHC-II | I-A/I-E | APC-Fire750 |  | 780/60 | 400 | Biolegend |
| CD117 | 2B8 | BV421 | Violet  (405 nm) | 450/50 | 50 | Biolegend |
| Viability dye | NA | BV510 |  | 525/50 | 100 | Thermo Fisher |
| CD4 | RM4-5 | BV570 |  | 586/15 | 400 | Biolegend |
| Ly6G | 1A8 | BV605 |  | 605/40 | 200 | BD Biosciences |
| CD19 | 6D5 | BV650 |  | 677/20 | 200 | Biolegend |
| Siglec-F | E50-2440 | BV711 |  | 710/50 | 200 | BD Biosciences |
| CD71 | C2 | BV750 |  | 750/30 | 100 | BD Biosciences |
| CD64 | X54-5/71 | BV786 |  | 810/40 | 100 | BD Biosciences |
| Nestin | 307501 | PE | Yellow-Green (561 nm) | 586/15 | 400 | Thermo Fisher |
| NKp46 | 29A1.4 | PE-Dazzle 594 |  | 610/20 | 50 | Biolegend |
| F4/80 | BM8 | PE-Cy5 |  | 670/30 | 100 | Biolegend |
| CD8 | 53-6.7 | PE-Cy5.5 |  | 710/50 | 400 | Thermo Fisher |
| CD11c | N418 | PE-Cy7 |  | 780/60 | 200 | Biolegend |
| TruStain FcX (CD16/32) | 93 | NA | NA | NA | 100 | Biolegend |
| Brilliant Stain Buffer Plus | N/A | N/A | N/A | N/A | 5 | BD Biosciences |
